# Calcium signaling through a Transient Receptor Channel is important for *Toxoplasma gondii* growth

**DOI:** 10.1101/2020.10.09.332593

**Authors:** Karla M. Márquez-Nogueras, Nathan M. Chasen, Myriam A. Hortua Triana, Ivana Y. Kuo, Silvia N.J. Moreno

## Abstract

Transient Receptor Potential (TRP) channels participate in ion calcium (Ca^2+^) influx and intracellular Ca^2+^ release. TRP channels have not been studied in *Toxoplasma gondii* or any other Apicomplexan parasite. We characterized a protein predicted to possess a TRP domain (TgTRPPL-2) and determined its role in Ca^2+^ signaling in *T. gondii*, the causative agent of toxoplasmosis. TgTRPPL-2 localized to the plasma membrane and the endoplasmic reticulum of *T. gondii*. The *ΔTgTRPPL-2* mutant was defective in growth and Ca^2+^ influx. Heterologous expression of TgTRPPL-2 in HEK-3KO cells allowed its functional characterization. Patching of ER-nuclear membranes demonstrated that TgTRPPL-2 is a non-selective cation channel that conducts Ca^2+^. Pharmacological blockers of TgTRPPL-2 inhibited Ca^2+^ influx and parasite growth. This is the first report of an Apicomplexan channel that conducts Ca^2+^ and initiates the Ca^2+^ signaling cascade that culminates in the stimulation of motility, invasion and egress. TgTRPPL-2 is a potential target for combating toxoplasmosis.

## INTRODUCTION

Ca^2+^ signaling is universal and forms part of the signaling pathways that activate or modulate a variety of physiological responses such as gene transcription, muscle contraction, cell differentiation and proliferation [1]. Ca^2+^ signals can be generated through the opening of ion channels that allow the downward flow of Ca^2+^ from either outside the cell or from intracellular stores like the endoplasmic reticulum [2].

*Toxoplasma gondii* is an intracellular parasite from the Apicomplexan phylum, that causes toxoplasmosis in humans [3]. In immunocompromised individuals, infection with *T. gondii* may lead to severe complications like encephalitis, myocarditis and death [4]. The *T. gondii* tachyzoite engages in a lytic cycle directly responsible for the pathogenicity of the infection as it results in lysis of host cells [5]. The lytic cycle consists of active invasion of host cells, replication inside a parasitophorous vacuole, followed by egress to search for a new host cell to invade. Ca^2+^ signals resulting from Ca^2+^ influx or from intracellular release, trigger a signaling cascade in the parasite that culminates in the stimulation of essential features of its lytic cycle, like motility, invasion, egress and secretion of proteins essential for attachment to the host cell [6, 7].

Previously we demonstrated the presence of a Ca^2+^ influx activity at the plasma membrane of *T. gondii* tachyzoites that was functional in extracellular tachyzoites [8] and intracellular replicating parasites [9]. The application of voltage operated Ca^2+^ channel blockers such as nifedipine inhibited ~ 80% of Ca^2+^ influx, and the residual Ca^2+^ entry activity suggested the potential existence of more than one channel at the plasma membrane of *T. gondii* [8]. The molecular entity of these channels has remained elusive.

Transient Receptor Potential (TRP) channels are a large family of cation permeable channels grouped into seven subfamilies based upon their gene sequence [10]. TRP channels can be activated by a multitude of stimuli and are involved in a wide range of cellular functions [11]. Most TRP channels are permeable to Ca^2+^ and all of them are permeable to monovalent cations [11]. Some TRP channels can participate in Ca^2+^ influx as well as Ca^2+^ release from intracellular stores [12, 13]. Mutations in these molecules are associated with a diverse set of diseases, due to their wide distribution in various tissues and their roles in pathological conditions, like cancer and renal diseases, making these channels important therapeutic targets [14]. The polycystin TRP (TRPP) subfamily of proteins are implicated in Autosomal Dominant Polycystic Kidney Disease (ADPKD) [15].

Predicted protein sequences with TRP domains have been found in most parasitic protozoa, although in lower numbers and types than in other organisms [16]. This could be the result of the evolutionary distance between the species studied, or because of loss of specific functions resulting from evolution of the parasitic lifestyle [16]. A genome analysis of a number of pathogenic protozoan parasites [17] searching for genes with homology to mammalian Ca^2+^ channels identified two *T. gondii* hypothetical genes (TgGT1_247370 and TgGT1_310560) with homologous regions to the TRPP family [18]. We termed these genes *TgTRPPL-1* and *TgTRPPL-2* (TRPP <u>L</u>ike) respectively. Previous work from our laboratory, localized the gene product of *TgTRPPL-1* to the ER with high-affinity tags due to its low level of expression [19].

In the current study we characterize TgTRPPL-2 in *T. gondii*, which represents the first TRP cation channel studied in any Apicomplexan parasite. Using reverse genetic approaches, we determine the role of TgTRPPL-2 in the lytic cycle of *T. gondii*. We also characterize the electrophysiological features of TgTRPPL-2 and its role in Ca^2+^ influx and, interestingly, find that pharmacological agents that block the activity of TgTRPPL-2 also inhibit Ca^2+^ influx in the parasite and parasite growth. TgTRPPL-2 emerges as one of the molecular entities involved in initiating Ca^2+^ signals in *T. gondii*.

## RESULTS

### TgTRPPL-2 localizes to the plasma membrane and the endoplasmic reticulum

Hypothetical proteins encoded by two genes in the *T. gondii’s* genome are annotated as proteins possessing Polycystic Kidney Disease (PKD) domains, which are characteristic of the subfamily P, or polycystin, TRP channels. Mammalian TRP-P channels contain 6 transmembrane domains with an extracellular loop between the first and second transmembrane domain [20]. We termed the *T. gondii* proteins TgTRPPL-1 and TgTRPPL-2. BLAST analysis to compare the sequences of the mammalian *PKD2* and *TgTRPPL-2* revealed low sequence homology (21.7%), even within the PKD Domains. The *TgGT1_310560* gene predicts the expression of a protein of 2,191 amino acids with an apparent molecular weight of 237 kDa and 14 transmembrane domains. The predicted topology [21] shows an extracellular loop between the first and second transmembrane domain, which is characteristic of PKD channels (Fig. 1A, *TgGT1_310560 cartoon*). While our BLAST analysis showed low sequence homology, HHPred, which searches for homologs based on protein sequence and secondary structure [22] revealed that TgTRPPL-2 shows significant homology with mammalian PKD2 (Table S1).

**Figure 1.**
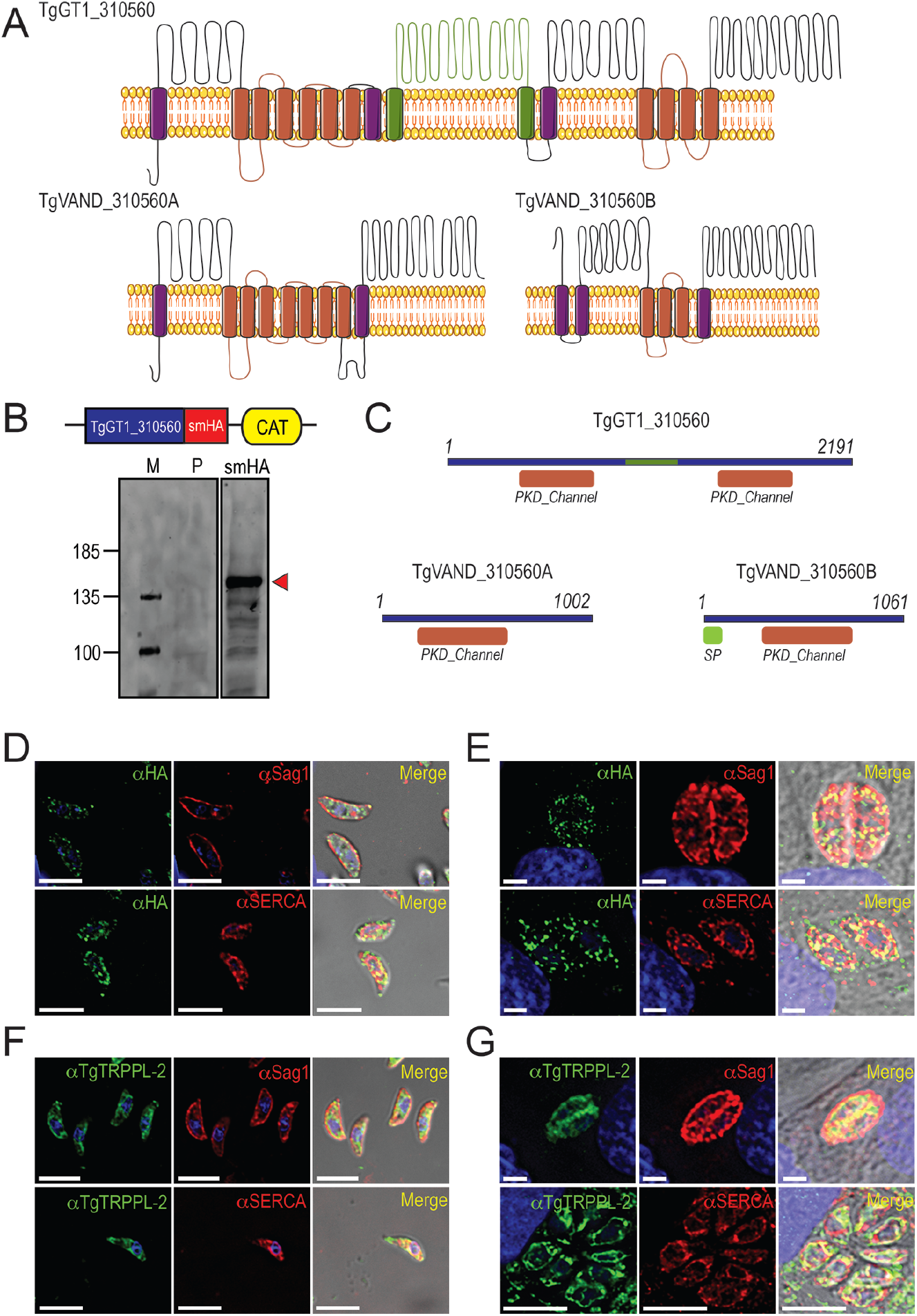
TgTRPPL-2 localizes to the plasma membrane (PM) and ER of *T. gondii*. **A.** Predicted topology for TgTRPPL-2 in GT1 and VAND strains. Model was generated with the Protter application [21]. The PKD Domain is highlighted in orange. The domain used to generate antibodies is highlighted in green. **B.** Schematic representation of C-terminal tagging of TgTRPPL-2 in TatiΔKu80 parasites and western blots of TgTRPPL-2-smHA membranes using αHA (1:1,000) showing a major band at approximately 150 kDa (*red arrowhead*). **C.** Schematic representation of the InterPro Domain annotation of TgTRPPL-2 in GT1 and VAND strains. **D.** IFAs of extracellular tachyzoites using αHA antibody show vesicular staining close to the PM and intracellular. Co-localization with αSAG1 and αSERCA show partial co-localization with both markers. **E.** IFAs of intracellular tachyzoites with αHA (1:100), αSag1 (1:1,000) and αSERCA (1:1,000) showing co-localization at the PM and ER. **F-G.** IFAs of extracellular and intracellular tachyzoites respectively with αTRPPL-2 (1:1,000) showing labeling of the protein at the periphery, co-localized with αSAG1 (1:1,000) and ER co-localized with αTgSERCA (1:1,000).

To investigate the localization of TgTRPPL-2, we introduced the high-affinity tag smHA [19] at the 3’ terminus of the *TgTRPPL-2* locus and isolated TgTRPPL-2-smHA cell clones. Carboxy-terminus tagging was done in the parental line RHTatiΔku80 (*TatiΔku80*) which favors homologous recombination [23]. Correct incorporation of the tag in the TgTRPPL-2-smHA line was validated by PCR (Fig. S1A) and western blot analysis using anti-HA antibodies (Fig. 1B). A band of approximately ~150 kDa was observed in lysates of TgTRPPL-2-smHA tachyzoites, which is nearly 87 kDa smaller (Fig. 1B) than the predicted size of 237 kDa without taking into account the smHA tag (~39 kDa).

Interestingly, a recent release of ToxoDBv.45 presents additional models for the *TgGT1_310560* gene from different *T. gondii* strains. The gene model for *TgVAND_310560* shows two fragments, *TgVAND_310560A,* which predicts a protein with 9 TMD and a size of ~107.97 kDa and *TgVAND_310560B*, which predicts a protein with 6 TMD and a size of ~116.6 kDa. Sequence alignments of the *TgGT1_310560* gene with the gene models for the VAND strain (A and B) shows 98% homology between them (Fig. 1C). The *T. gondii* VAND strain is an isolate from South America and belongs to the hypervirulent Type I group, as the GT1 and RH strains. According to the gene model of the VAND strain, the predicted MW of the TgVAND_310560B protein would be similar to the band size observed in our western blots (116.6 + 39) with a predicted topology of 6 TMD, in closer agreement with typical TRP channels [24, 25].

To further demonstrate that the protein band observed in the western blot analysis corresponded to the tagged TgTRPPL-2 gene, we performed immunoprecipitations with anti-HA of lysates from the TgTRPPL-2-smHA cells. The immunoprecipitated samples were run in a PAGE gel. The ~150 kDa band was excised and analyzed by mass spectrometry (Fig. S1B, *red box*)(Table S2). Approximately ~700 amino acids toward the C-terminus domain of the fusion protein were covered by the peptides, corresponding to a 66% coverage. Comparison of the coverage with the TgVAND_310560B predicted protein, revealed that approximately 66% of the sequence was recovered by the mass spectrometry analysis. This result indicates that the TgGT1_310560 is likely cleaved, a characteristic in common with other TRP channels [26].

We next investigated the cellular localization of TgTRPPL-2. Immunofluorescence analysis (IFAs) of extracellular and intracellular parasites showed that TgTRPPL-2-smHA apparently localizes to peripheral vesicles close to the plasma membrane (PM) and to the endoplasmic reticulum (ER) (Fig. 1D-E). Some co-localization with the plasma membrane surface antigen (Sag1) and the sarco-endoplasmic reticulum Ca^2+^-ATPase (TgSERCA) (ER marker) was observed (Fig. 1D-E). However, considering the low-level of expression of TgTRPPL-2, it was difficult to draw definitive conclusions about its localization.

We next produced polyclonal antibodies against a fragment peptide of TgTRPPL-2, which is indicated in Fig. 1A (*highlighted in green*). The peptide was expressed in bacteria, purified and used for immunization of mice. Mouse serum was isolated and affinity purified prior to its use for IFA. The localization at the periphery of extracellular tachyzoites was further confirmed by co-localization with αSAG1 (Fig. 1F). Extracellular tachyzoites showed intracellular staining that co-localized with TgSERCA (Fig. 1F) supporting ER localization. Additionally, IFA of intracellular tachyzoites showed that TgTRPPL-2 co-localizes with αSAG1 and αSERCA (Fig.1G).

In summary, TgTRPPL-2 is expressed in *T. gondii* tachyzoites, it is likely post-translationally cleaved, and localizes to the PM and the ER.

### TgTRPPL-2 is important for growth, invasion and egress of Toxoplasma gondii

With the aim of investigating the physiological role of TgTRPPL-2 in *T. gondii*, we generated a *ΔTgTRPPL-2* mutant using the CRISPR-Cas9 approach to disrupt the transcription of *TgTRPPL-2* by inserting a dihydrofolate reductase-thymidylate synthase (DHFR) cassette in the *TgTRPPL-2* genomic locus (Fig. 2A). Genetic controls for the insertion were done by PCR, and qPCR showed a significant decrease in the levels of *TgTRPPL-2* transcripts (Fig. 2B).

**Figure 2.**
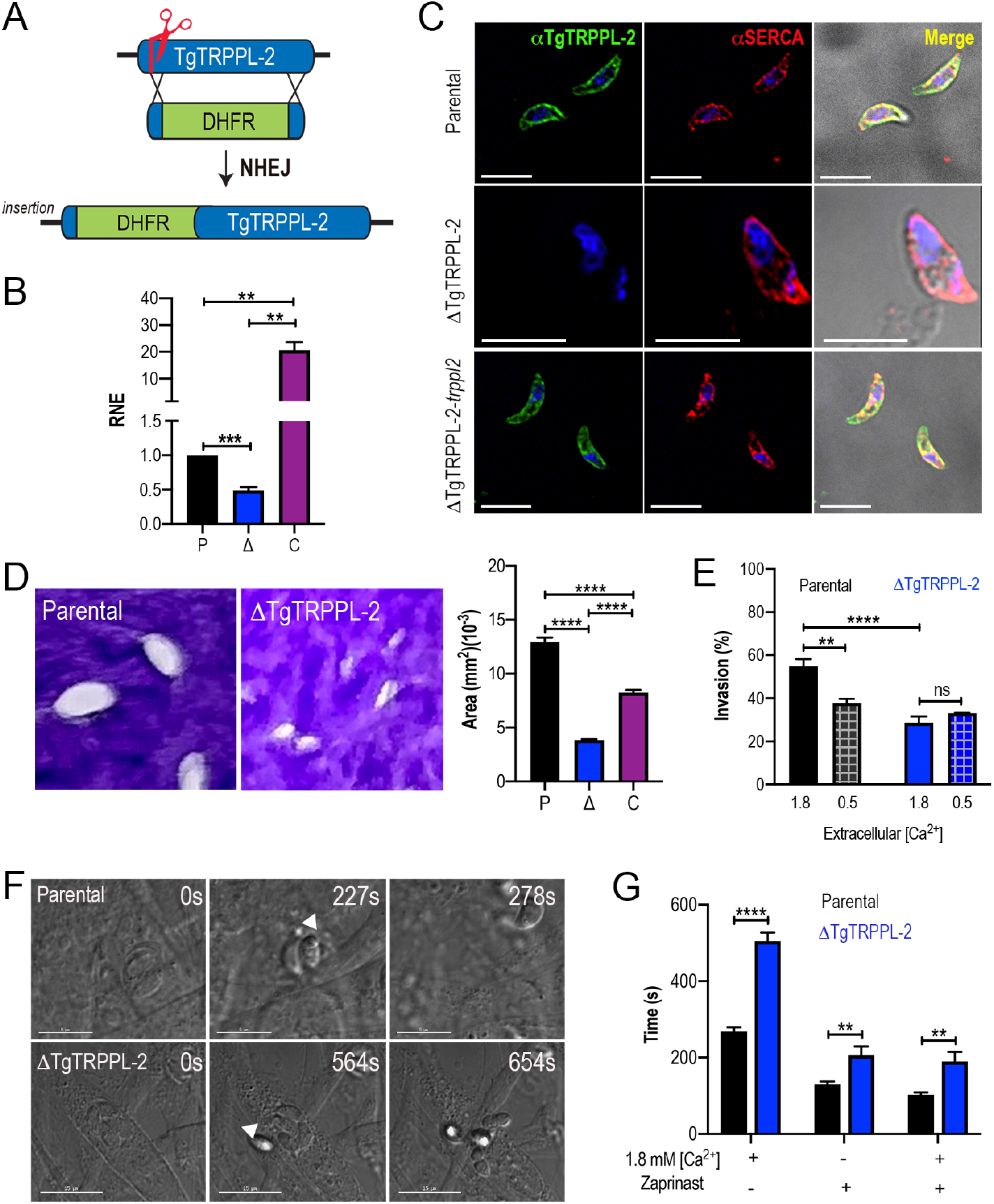
The role of TgTRPPL-2 in *T. gondii* growth. Schematic representation of the generation of *ΔTgTRPPL-2* in the *T. gondii* RH strain. **B.** qPCR of total RNA from *ΔTgTRPPL-2*, *ΔTgTRPPL-2*-*trppl2* and parental strains using primers upstream and downstream of the insertion site of the DHFR cassette. **C.** IFAs of extracellular parasites showing PM labeling with αTgTRPPL2 (1:1,000) and co-localization with αSERCA (1:1,000). **D**. Plaque assays of parental (P), *ΔTgTRPPL-2* (Δ) and *ΔTgTRPPL-2*-*trppl2* (C) parasites. Quantification of plaque sizes from three independent biological experiments using Student’s *t*-test. \*\*\*\**p* < 0.0001. **E.** Red green assays of parental, *ΔTgTRPPL-2* and *ΔTgTRPPL-2-trppl2* cells quantifying invaded and attached intracellular parasites. Assays were done at two concentrations of extracellular Ca^2+^: 0.5 and 1.8 mM. Values are means ± SEM. ** *p* < 0.001, **** *p* < 0.0001. **F.** Time to egress stimulated by saponin/Ca^2+^ at 1.8 mM extracellular Ca^2+^ of both parental or the *ΔTgTRPPL-2* mutant. **G.** Statistical analysis of average egress time stimulated by saponin or Zaprinast in the presence or absence of extracellular Ca^2+^. Analysis was performed from three independent biological replicates using Student’s t-test. Values are means ± SEM, ** *p* < 0.003, **** *p* < 0.0001. *Black bars* represent parental strain, *blue bar*s represent ΔTgTRPPL-2.

The *ΔTgTRPPL-2* mutant was complemented with Cosmid PSBLZ13 [27] that contains the whole genomic locus of the *TgTRPPL-2* gene to generate the cell line *ΔTgTRPPL-2*-*trppl2*. Controls for the expression of *TgTRPPL-2* were done by qPCR and by IFAs, which further confirmed the identity of the tagged protein, as it was not expressed in the *ΔTgTRPPL-2* mutant and was present in the complemented line *ΔTgTRPPL-2*-*trppl2* (Fig. 2B-C). Further validation of the absence of expression of TgTRPP2-L and its complementation is shown in Figs S2A-B with additional IFA images and western blots of *ΔTgTRPPL-2* and *ΔTgTRPPL-2*-*trppl2* parasites (Fig S2B).

We next evaluated if the expression of *TgTRPPL-2* would impact *T. gondii* growth by plaque assays, in which the parasite engages in repetitive cycles of invasion, replication, and egress causing host cell lysis and formation of plaques observed as white spots by staining with crystal violet. The *ΔTgTRPPL-2* mutant formed smaller plaques compared to its parental counterpart indicating a growth defect (Fig. 2D). This growth defect was partially restored in the complemented cell line (Fig. 2D). We reasoned that the overexpression of *TgTRPPL-2* in the *ΔTgTRPPL-2*-*trppl2* mutant may affect parasite fitness masking the rescue effect.

To determine which step of the lytic cycle was affected we performed invasion and egress assays. For invasion we used the red-green assay [28] under two extracellular Ca^2+^ concentrations (1.8 and 0.5 mM). Quantification of invasion in the presence of 1.8 mM Ca^2+^ showed a lower invasion rate for the *ΔTgTRPPL-2* (Fig. 2E). Reducing the extracellular concentration of Ca^2+^ to 0.5 mM resulted in a reduced rate of invasion by the parental line, which was similar to the invasion rate of the *ΔTgTRPPL-2* mutant. This result demonstrated that TgTRPPL-2 is important for invasion at higher concentrations of extracellular Ca^2+^.

Egress of intracellular tachyzoites can be triggered by permeabilizing infected host cells with saponin in the presence of a buffer containing 1.8 mM of extracellular Ca^2+^. Under these conditions egress of the *ΔTgTRPPL-2* mutant was slower than egress of the parental strain (Fig. 2F). Additionally, when egress was stimulated by Zaprinast, which increases cytosolic Ca^2+^, the *ΔTgTRPPL-2* mutant also took longer to egress (Fig. 2G). For both assays tested, the *ΔTgTRPPL-2* mutant took twice as long as the parental line.

In summary, disruption of the TgTRPPL-2 locus negatively impacted two important steps of the *T. gondii* lytic cycle, invasion and egress, which impacted parasite growth.

### The role TgTRPPL-2 in Ca^2+^ influx

We previously showed that *T. gondii* tachyzoites allow influx of Ca^2+^ when exposed to 2 mM extracellular Ca^2+^ [8]. To determine the role of TgTRPPL-2 in this pathway, we loaded *ΔTgTRPPL-2* parasites with Fura-2AM to study intracellular Ca^2+^ changes after exposing them to 1.8 mM extracellular Ca^2+^ (Fig. 3A). The resting cytosolic Ca^2+^ concentration of *ΔTgTRPPL-2* mutant was around 75 nM which is similar to the resting concentration of parental cells (~70-100 nM). Adding 1.8 mM Ca^2+^ to the extracellular buffer caused an increase in cytosolic Ca^2+^ in both the parental strain and the *ΔTgTRPPL-2* mutant (Fig. 3B). However, Ca^2+^ influx of the *ΔTgTRPPL-2* mutant was significantly lower (Fig. 3C) and was decreased by almost 50%. The *ΔTgTRPPL-2*-*trppl2* complemented mutant, however, regained the Ca^2+^ influx activity and it even showed higher Ca^2+^ influx than parental cells, consistent with the higher expression of *TgTRPPL-2* shown by qPCR (Fig. 2B). The reduction of Ca^2+^ influx was further confirmed when adding 1 mM extracellular Ca^2+^ to the *ΔTgTRPPL-2* mutant (Fig. S3A).

**Figure 3.**
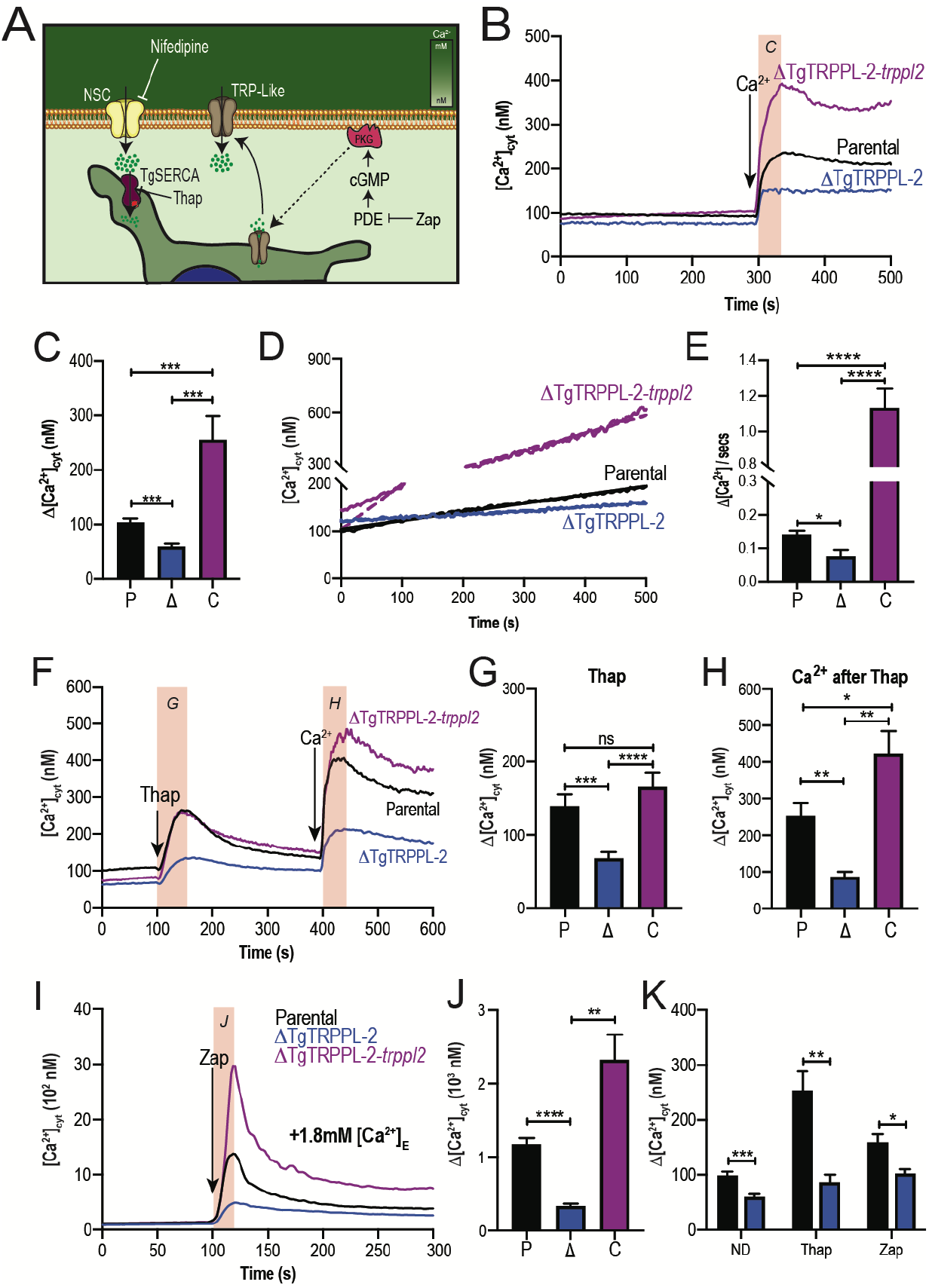
The role of TgTRPPL-2 in PM Ca^2+^ influx. **A.** Scheme showing the mechanism of Ca^2+^ influx and how cytosolic Ca^2+^ may activate the PM channel (Ca^2+^-activated Calcium Entry). *NSC, Nifedipine-Sensitive Channel; PKG, Protein Kinase G; PDE, phosphodiesterase; Thap, Thapsigargin.* **B.** Cytosolic Ca^2+^ measurements of Fura-2 loaded tachyzoites of the parental (P), *ΔTgTRPPL-2* (Δ) and *ΔTgTRPPL-2*-*trppl2* (C) lines. The buffer contains 100 μM EGTA and at 300 sec, 1.8 mM Ca^2+^ was added to the suspension. The *orange box* indicates the area used for the quantification presented in *C*. **C.** Quantification and statistical analysis of the change in cytosolic Ca^2+^ during the first 20 s after addition of extracellular Ca^2+^. *** *p* < 0.0002. **D.** Cytosolic Ca^2+^ increase of parasites pre-incubated with 1.8 mM Ca^2+^. **E.** Quantification and statistical analysis of the slope from D. **** *p* < 0.0001. **F.** Ca^2+^ efflux after adding Thap (1 *μ*M) followed by Ca^2+^ influx after the addition of 1.8 mM extracellular Ca^2+^. The *orange boxes* indicate the area used for the quantification presented in *G* and *H*. **G.** Quantification and statistical analysis of the change in cytosolic Ca^2+^ 50 s after the addition of Thap *(Thap)* and **(H)** 20 s after the addition of 1.8 mM of Ca^2+^ *(Ca*^2+^ *after Thap)*. *** *p* < 0.0008, **** *p* <0.0001. **G.** Cytosolic Ca^2+^ increase stimulated by Zaprinast (100 *μ*M) in the presence of 1.8 mM extracellular Ca^2+^. **J.** Quantification and statistical analysis of cytosolic Ca^2+^ increase during the first 15 s after adding Zaprinast (100 *μ*M) (*Orange box*, in I). ** *p* < 0.001, **** *p* < 0.0001. **K.** Quantification and statistical analysis of Ca^2+^ influx during the 20 s after adding Ca^2+^ without additions (ND) or after adding Thap or Zap. * p < 0.02, ** p< 0.005, *** p< 0.0008. Statistical analysis for all experiments were done from at least three independent trials using student’s t-test.

When *T. gondii* tachyzoites are suspended in a high Ca^2+^ buffer from the beginning of the experiment there is a slow constitutive influx of Ca^2+^, that we attribute to leakage through a PM channel (Fig. 3D, *parental black tracing*). Interestingly, this leakage activity was significantly reduced in the *ΔTgTRPPL-2* mutant (Fig. 3D-E, *blue tracing and bar*), supporting a role of TgTRPPL-2 in constitutive Ca^2+^ influx at the PM. Additional evidence is provided by the enhanced Ca^2+^ leakage observed with the *ΔTgTRPPL-2*-*trppl2* complemented mutant (Fig. 3D-E, *purple tracing and bar*). The high Ca^2+^ leakage and Ca^2+^ influx observed with the *ΔTgTRPPL-2*-*trppl2* mutant may affect parasite fitness and would explain the partial growth recovery observed in the complemented mutant.

Ca^2+^ channels may also be modulated by Ca^2+^ itself [29]. We previously showed that a cytosolic [Ca^2+^] increase may activate Ca^2+^ influx at the PM (Ca^2+^ activated-Ca^2+^ entry) [8]. We next investigated if the Ca^2+^ activated-Ca^2+^ entry (CACE) activity was due to the functioning of TgTRPPL-2 at the PM. We added thapsigargin (thap) to tachyzoites in suspension (Fig. 3F), which results in a cytosolic Ca^2+^ increase due to inhibition of the SERCA-Ca^2+^-ATPase (SERCA) resulting in uncompensated Ca^2+^ efflux into the cytosol. This elevated cytosolic Ca^2+^ stimulates further Ca^2+^ influx at the PM, which we assessed as an increase in cytosolic Ca^2+^ following addition of high Ca^2+^ to the buffer (Fig 3F, *black tracing*). Note that the Δ[Ca^2+^]_cyt_ shown in Fig. 3H, *black column*, is almost 2.5 times higher than the Δ[Ca^2+^]_cyt_ observed without previous addition of thap (Fig. 3C, *black column*). This CACE activity was absent in the *ΔTgTRPPL-2* mutant (Fig. 3F, *blue tracing*) but was restored in the *ΔTgTRPPL-2*-*trppl2* complemented strain (Fig. 3F, *purple tracing*). Quantifications of the rate of Ca^2+^ increase after adding Ca^2+^ and statistical analyses are shown in Fig. 3G-H. Note that the *ΔTgTRPPL-2* mutant showed a reduced response to the addition of thap and also to the addition of Ca^2+^. Comparing the response to the addition of extracellular Ca^2+^ shown in Fig. 3H, *blue column*, the Δ[Ca^2+^]_cyt_ is similar to that measured directly without previous addition of thap (compare with the *blue column* in Fig 3C). This result points to a complete absence of the modulatory effect of cytosolic Ca^2+^ on the PM Ca^2+^ influx in the *ΔTgTRPPL-2* mutant, which is restored in the complemented mutant, *ΔTgTRPPL-2*-*trppl2*.

We next tested Zaprinast, which increases the levels of cGMP resulting in Ca^2+^ release from an unidentified store [30]. We previously determined that Ca^2+^ release was almost 2.5 times higher in the presence of extracellular Ca^2+^, as compared with that in the absence of extracellular Ca^2+^ [30]. We attribute this increase to stimulation of the PM Ca^2+^ channel by cytosolic Ca^2+^ (CACE). When testing this phenotype with the *ΔTgTRPPL-2* mutant, we observed that the increased response was absent (Fig. 3I-K). Also note that even the release of Ca^2+^ from intracellular stores by Zaprinast in the presence of low extracellular Ca^2+^ (~50 nM) was significantly decreased in the *ΔTgTRPPL-2* mutant (Fig. S3B-C). The modulatory action of elevated cytosolic Ca^2+^ in Ca^2^ influx was absent in the *ΔTgTRPPL-2* mutant (Fig. 3I-K).

Taken together, these results suggest a role for TgTRPPL-2 in Ca^2+^ influx at the PM. In addition, TgTRPPL-2 is modulated by cytosolic Ca^2+^ and is responsible for a constitutive PM Ca^2+^ influx pathway.

### TgTRPPL-2 is a cation conducting channel

With the aim of establishing that TgTRPPL-2 functions as a channel and is able to conduct Ca^2+^ we cloned the cDNA of the *TgTRPPL-2* gene into a mammalian expression vector (pCDNA3.1) for expression in human embryonic kidney 293 cells (HEK-3KO) [31]. These HEK cell line is genetically modified and the 3 isoforms of the inositol 1,4,5-trisphosphate receptor (IP_3_R) are deleted, to reduce background Ca^2+^ currents [31]. TgTRPPL-2 was localized to the ER of HEK cells as assessed by co-localization with a red fluorescent protein (RFP) targeted to the ER and compared with the mammalian homolog polycystin 2 (PC2) (Fig. 4A). Because of this, we isolated nuclear/ER membranes (Fig. 4B) for single channel patch clamp experiments and further characterization of the permeability properties of TgTRPPL-2 (Fig. 4C).

**Figure 4.**
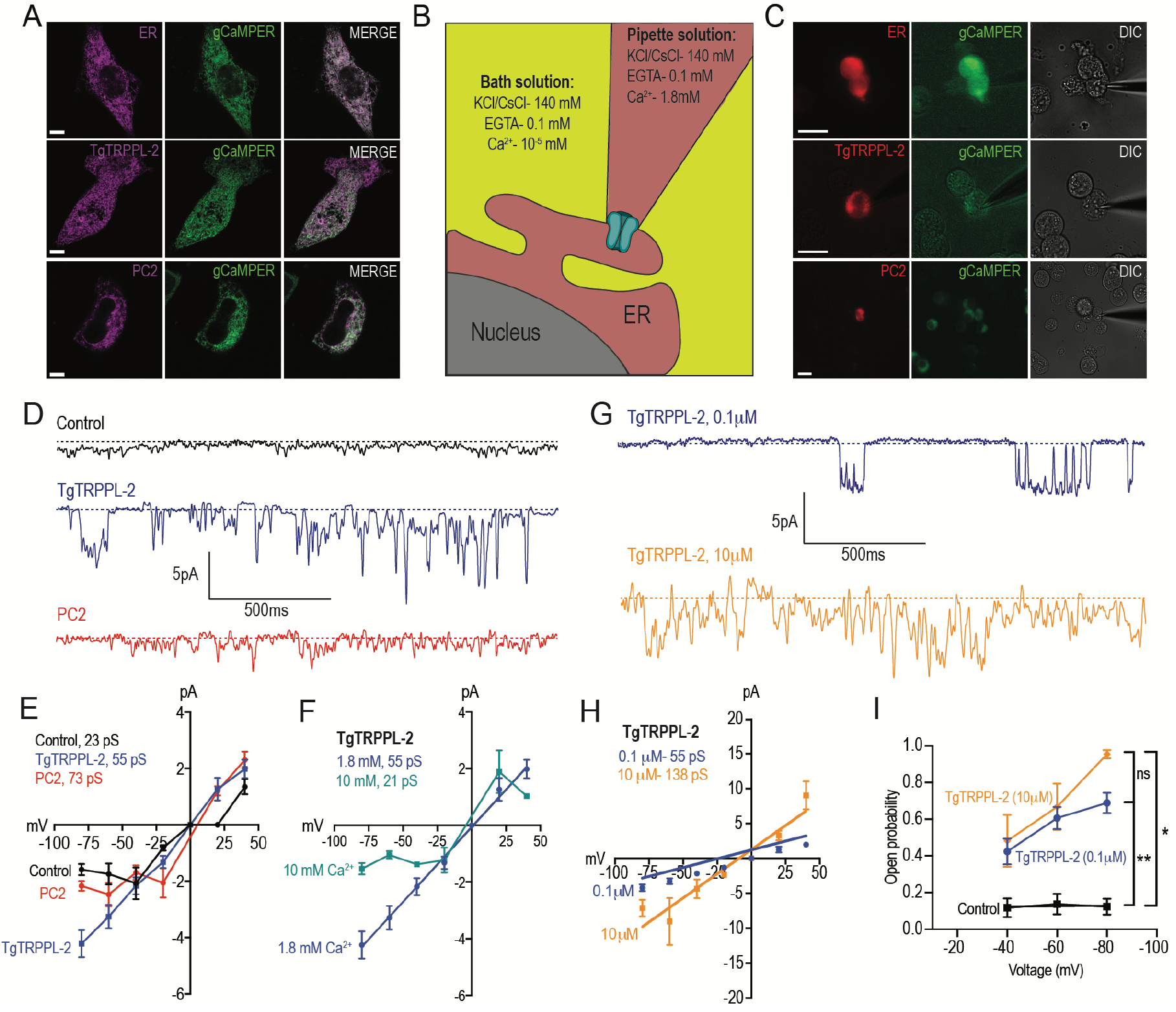
Characterization of TgTRPPL-2 expressed in HEK-3KO cells. **A.** Images of HEK-3KO cells expressing an ER-marker, PC2 or TgTRPPL-2 with the genetic calcium indicator gCAMPER. **B.** Schematic representation of nuclear-patch clamp. Ionic composition and concentration for bath and pipette solutions are shown. **C.** Patched nuclear-extract expressing ER-marker, PC2 and TgTRPPL-2 with the genetic calcium indicator gCaMPER. **D.** Representative tracing from control, *TgTRPPL-2* or *PC2* expressing cells showing the currents recorded in the presence of 1.8 mM luminal Ca^2+^ in a potassium chloride solution. Tracings represent approximately 2 sec from a 25 sec analysis and filtered using 45 kHz. **E.** Current-Voltage relationship comparing single-channel conductance of control, PC2 and TgTRPPL-2 expressing cells from *D*. *Inset*, calculated channel conductance for Control, TgTRPP-L 2 and PC2. **F.** Current-Voltage relationship comparing single-channel conductance of TgTRPPL-2 expressing cells at 1.8 and 10 mM [Ca^2+^] inside the pipette. *Inset*, calculated channel conductance for the conditions analyzed. **G.** Representative tracing of currents recorded from TgTRPPL-2 expressing cells using different concentration of [Ca^2+^] in the bath solution (Solution A vs. Solution B) (Table S4). Tracings represent approximately 2 sec from a 25 sec analysis and filtered using 45 kHz. **H.** Current-voltage relationship comparing single-channel conductance of TgTRPPL-2 expressing cells at 0.1 and 10 μM [Ca^2+^] in the bath solution. Inset, calculated channel conductance for the different [Ca^2+^]. **I.** Open probability of control and TgTRPPL-2 expressing cells in the presence of different [Ca^2+^] in the bath solution in comparison to the Control. * *p* < 0.01, ** *p* < 0.001.

In the presence of 1.8 mM of Ca^2+^ inside the patch pipette and 100 nM of Ca^2+^ in the bath solution (see scheme of Fig. 4B), the membranes isolated from control cells, held at −80 mV, showed very little activity and the conductance remained at less than 1.5 pA (Fig. 4D, *control tracing*). Some channel activity was observed after artificially depolarizing membranes (−80 to +20 mV) presumably due to opening of potassium channels. In comparison, when analyzing membranes isolated from cells expressing TgTRPPL-2, a significant increase in the open probability and current sizes was observed (Fig. 4D, *TgTRPPL-2 blue tracing*). The current-voltage relationship was linear and significantly different from the one from control cells (Fig. 4E, *blue vs. black line*).

We compared the activity of TgTRPPL-2 with that of the mammalian PKD channel PC2 in parallel experiments since PC2 has been well characterized in the literature. Activity of PC2 expressing cells displayed a voltage-dependent behavior, as the current-voltage relationship was not linear, with a conductance of ~73 pS (Fig. 4D-E, *red tracing*). Previous work has demonstrated that PC2 can be voltage dependent. Additionally, depending on the experimental design, the measured conductance for PC2 can be variable. Comparing our experimental approach to previous work, conductance for PC2 in a high Ca^2+^ solution is similar (~73 pS vs. ~97 pS). Although TgTRPPL-2 does not appear to be voltage dependent, conductance of the channel is similar to that of its mammalian homologue.

Ca^2+^ is able to modulate the activity of channels like TRP-P channels, some of which have an EF-hand motif at the C-terminus, and have been shown to be activated by Ca^2+^ [29, 32]. To determine whether Ca^2+^ is able to modulate the activity of TgTRPPL-2 we first varied the [Ca^2+^] inside the pipette (in equilibrium with the ER lumen). When the Ca^2+^ concentration was increased to 10 mM Ca^2+^ there was a significant inhibition of TgTRPPL-2 channel activity. With high Ca^2+^ concentration, the channel displayed voltage-dependent inhibition over the −75 to −25 mV range and conductance was significantly decreased (Fig. 4F). In the presence of 1.8 mM Ca^2+^ inside the pipette, TgTRPPL-2 had a conductance of ~55 pS, which is significantly higher than the conductance calculated for control membranes. The conductance decreased to ~21 pS when Ca^2+^ was increased to 10 mM.

Although no evidence for a conserved EF-hand motif was found in TgTRPPL-2 we checked for the potential modulation by cytosolic Ca^2+^. Increasing the concentration of Ca^2+^ in the bath solution from 100 nM to 10 μM (which would simulate changes in cytosolic Ca^2+^), enhanced channel activity from membranes expressing TgTRPPL-2 (Fig. 4G-H, *blue vs. gold line*). Interestingly, increasing the [Ca^2+^] only increased the open probability when the membrane was depolarized to −80 mV (Fig. 4I). However, increase of the [Ca^2+^] of the bath solution, increased the conductance of the channel almost 2.5x, suggesting modulation of the channel by Ca^2+^ itself. In conclusion, these data indicate that TgTRPPL-2 is able to conduct Ca^2+^ currents and is modulated by cytosolic Ca^2+^.

To distinguish whether TgTRPPL-2 is able to conduct cation currents and to determine if the activity measured could be the result of permeation of potassium, we replaced potassium with the non-permeable ion cesium [33, 34]. In the presence of 1.8 mM Ca^2+^ inside the pipette, in a cesium chloride solution, membranes from TgTRPPL-2- and PC2-expressing cells have a significantly higher activity than control cells (Fig. 5A). The current-voltage relationship is linear through different applied voltages and significantly different from that of control cells in potassium or cesium chloride solution (Fig. 5B). Although channel conductance is slightly higher in potassium chloride, it is not significantly different than the calculated conductance and open probability obtained in cesium chloride (Fig. 5B-C). However, when applying voltages higher than −40 mVs, the channel was open for longer times in the presence of cesium chloride vs potassium chloride (Fig. 5D, *green vs. blue line*). These results indicate that TgTRPPL-2 permeates Ca^2+^, however it can also conduct potassium, since channel conductance is slightly higher in the potassium chloride solution.

**Figure 5.**
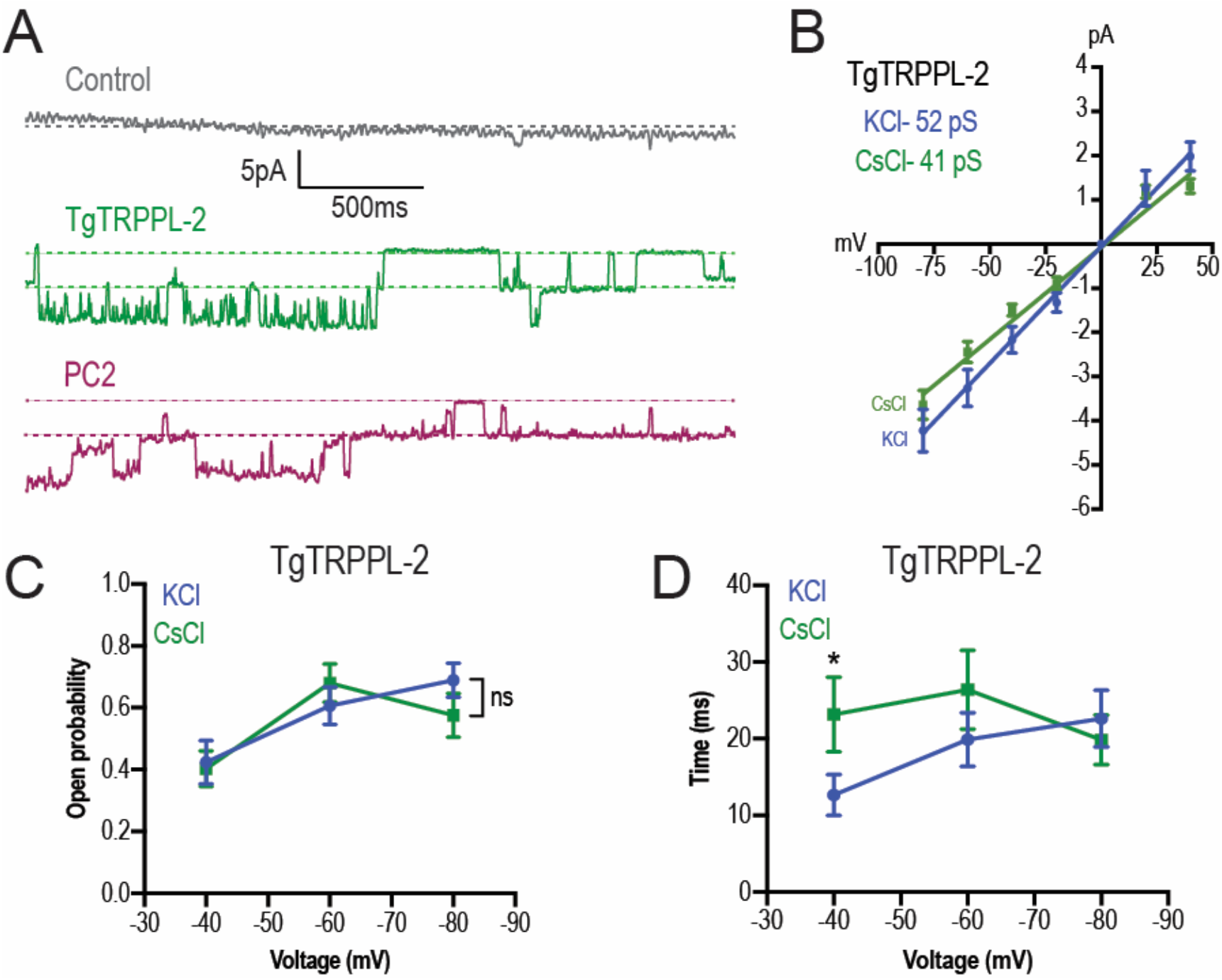
TgTRPPL-2 permeates Ca^2+^. **A.** Representative tracing of currents recorded at −80 mV in the presence of 1.8 mM Ca^2+^ inside the pipette (Solution D, Table S4) of nuclear extracts from Control, *TRPPL-2*- or *PC2*-expressing cells. Traces are a representation of 2 seconds of sampling from a total time of 25 seconds. **B.** Current-voltage relationship comparing single-channel conductance of TgTRPPL-2 cells in 1.8 mM in KCl (*blue*) or CsCl (*green*) buffer. *Inset*, Channel conductance of TgTRPPL-2 in the different conditions analyzed. **C.** Calculated open probability of TgTRPPL-2 expressing cells in the presence of 1.8 mM Ca^2+^ in a KCl (*blue*) or CsCl (*green*) buffer. **D.** Average time of channel opening of TgTRPPL-2 expressing cells in the presence of 1.8 mM Ca^2+^ in a KCl (*blue*) or CsCl (*green*). * *p* < 0.04. Values are means ± SEM,

We further demonstrated that TgTRPPL-2 is able to conduct Ca^2+^ by following Ca^2+^ changes of TgTRPPL-2-HEK-3KO or RFP-ER-HEK-3KO cells co-transfected with a genetic Ca^2+^ indicator, allowing to follow ER luminal Ca^2+^ changes and current activity simultaneously. Luminal Ca^2+^ changes were followed through one cycle of membrane depolarization from −80 mV to 40 mV (Fig. S4A). The fluorescence of the Ca^2+^ indicator decreased in the TgTRPPL-2-expressing cells with time, as voltage was applied. In both potassium as well as cesium chloride solutions at 1.8 mM Ca^2+^ we observed that the fluorescence decrease was significantly larger when the HEK-3KO cells expressed *TgTRPPL-2* (Fig. S4B-C vs. D-E). The slope for the fluorescence decrease appeared higher in the cesium chloride solution than in the potassium solution, although was quite variable (Fig. S4F-G). In summary, the observed decrease in the fluorescence of the Ca^2+^ indicator supports the Ca^2+^ permeation activity of TgTRPPL-2, which agrees with the single channel conductance measurements.

### Inhibition of TgTRPPL2 by TRP Channel Inhibitors

Previously we demonstrated Ca^2+^ influx in *T. gondii* and its inhibition by L-type voltage gated Ca^2+^ channel blockers like nifedipine (Fig. 6A) [8]. Taking into account that TgTRPPL-2 is a cation permeable channel and localizes to the PM, we next investigated if the residual Ca^2+^ influx activity observed with the *ΔTgTRPPL-2* mutant could be blocked with nifedipine. Interestingly, pre-incubation of the *ΔTgTRPPL-2* parasites with nifedipine showed that the initial cytosolic Ca^2+^ was elevated and it was around ~400 nM (compared to 100 nM of the parental strain under identical conditions) in Ca^2+^-free buffer (Fig. 6B). Further addition of extracellular Ca^2+^ did not result in Ca^2+^ influx and only a slow steady cytosolic Ca^2+^ increase was observed (Fig. 6B). We attribute the higher cytosolic Ca^2+^ in the presence of nifedipine to leakage of stored Ca^2+^ into the cytosol as the cell is trying to compensate for the complete absence of Ca^2+^ influx. However, the absence of Ca^2+^ influx after adding extracellular Ca^2+^ in the presence of nifedipine, points to a role for TgTRPPL-2 in Ca^2+^ influx.

**Figure 6.**
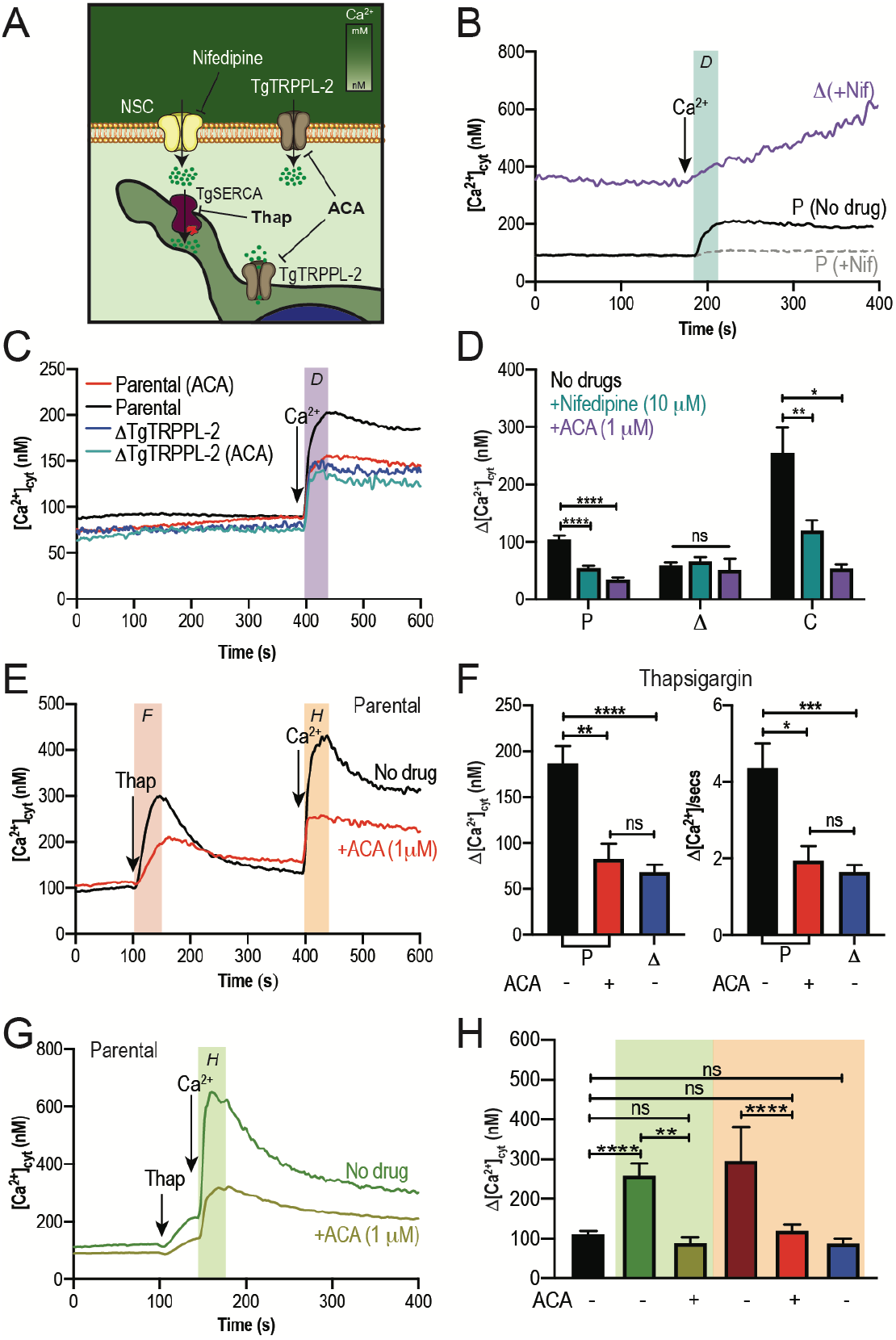
Regulation of TgTRPPL-2 by Ca^2+^ and inhibition by TRP inhibitors. **A.** Scheme showing TgTRPPL-2 at the PM and ER. **B**. Cytosolic Ca^2+^ measurements of Fura2 loaded tachyzoites pre-incubated with 10 *μ*M nifedipine. 1.8 mM Ca^2+^ was added where indicated. The *blue box* indicates the area used for the quantification presented in D. **C.** Cytosolic Ca^2+^ measurements of suspensions of parental and *ΔTgTRPPL-2* parasites pre-incubated for 3 min with ACA (1 *μ*M). 1.8 mM Ca^2+^ was added where indicated. The *purple box* shows the area used for the quantifications presented in D. **D.** Change in cytosolic Ca^2+^ during the first 20 s after addition of Ca^2+^ in the presence of 10 *μ*M of nifedipine or 1 *μ*M ACA. * p<0.01, ** p<0.003, **** p<0.0001. **E.** Cytosolic Ca^2+^ increase after adding Thap (1 *μ*M) to a suspension of tachyzoites. The red line shows a similar experiment but the cells were pre-incubated with ACA for 3 min. The *pink and orange boxes* show the areas used for the quantifications presented in F. **F.** Quantification and statistical analysis of the ΔCa^2+^ and slope 50 s after the addition of Thap in the presence or absence of ACA in parental and the *ΔTgTRPPL-2* mutant. *p* < 0.01, ** *p* < 0.003 *** *p* < 0.0003. **G.** Stimulation of Ca^2+^ influx by pre-addition of thap in the presence or absence of 1 *μ*M ACA. The green box shows the area used for the quantifications presented in H. **H.** Quantification of change of cytosolic Ca^2+^ 20 s after the addition of 1.8 mM Ca^2+^ following the addition of Thap under different conditions. ** *p* < 0.001, **** *p* < 0.00001. The statistical analysis for all experiments were done from at least three independent trials using student’s t-test. Values are means ± SEM.

We next tested the effect of anthranilic acid (ACA), a wide spectrum TRP channel inhibitor [35], on Ca^2+^ influx of both the parental control and the *ΔTgTRPPL-2* mutant (Fig. 6C). ACA inhibited Ca^2+^ influx by 40-50% of the parental cell line (Fig. 6C, *black vs. red tracing*). However, preincubation of *ΔTgTRPPL-2* tachyzoites with ACA, did not further reduce Ca^2+^ influx (Fig. 6C, *dark blue vs. light blue tracings*). Sensitivity to both nifedipine and ACA was restored in the complemented cell line (Fig. 6D). These results point to TgTRPPL-2 as a PM channel that conducts Ca^2+^, and it is relevant for its influx from the extracellular milieu, and is sensitive to TRP channel inhibitors.

### The Role of TRPPL-2 as a Ca^2+^ leak channel at the ER membrane

The dual localization of TgTRPPL-2 at the PM and ER suggests the potential function of the channel at both locations (Fig. 1D). Inhibition of TgSERCA with thap results in Ca^2+^ efflux into the cytosol [36]. We observed that incubation of tachyzoites of the parental strain with ACA significantly decreased the efflux of Ca^2+^ caused by thap (Fig. 6 E-F, *black line and bar vs. red line and bar*). The ACA-inhibited ER Ca^2+^ efflux rate was comparable to the decreased efflux rate triggered by thap in the *ΔTgTRPPL-2* mutant (Fig. 6F, *blue bar versus red bar and* Fig. 3D). In addition, Ca^2+^-activated-Ca^2+^ entry, evaluated by adding 1.8 mM of extracellular Ca^2+^ 50 s after stimulating efflux with thap, was inhibited by ACA (Fig. 6G, *green versus gold line*). Ca^2+^ influx after thap was reduced to basal Ca^2+^ influx (without pre-addition of thap) for both parental and *ΔTgTRPPL-2* mutant. Note that Ca^2+^ influx in *ΔTgTRPPL-2* tachyzoites after stimulus by thap, and Ca^2+^ influx in the parental strain without any stimulus are similar because the modulation of Ca^2+^ influx by cytosolic Ca^2+^ is lost in the *ΔTgTRPPL-2* parasites (Fig. 6H, *black versus blue bars*).

In conclusion, ACA inhibited both efflux of Ca^2+^ from the ER as well as Ca^2+^-induced-Ca^2+^ entry. This led us to propose that TgTRPPL-2, in addition to mediate Ca^2+^ influx at the plasma membrane may also mediate Ca^2+^ leakage from the ER, a pathway sensitive to the TRP-channel inhibitor ACA.

To further validate the specificity of ACA for the inhibition of TgTRPPL-2, we tested this inhibitor and a second broad spectrum TRP channel inhibitor, benzamil, against single channel conductance. Channel activity of TgTRPPL-2 was significantly decreased by both ACA and benzamil (Fig. 7A-B). ACA diminished the amplitude of the channel by reducing the probability and time that the channel remained open (Fig. 7C, E). Conductance of the channel was reduced to almost half in the presence of ACA (Fig. 7B), which correlates with the inhibition of Ca^2+^ entry in *T. gondii.* In comparison to ACA, benzamil only reduced the open probability of TgTRPPL-2 but not the length of time the channel was open (Fig. 7D-E)). The conductance of TgTRPPL-2 was reduced to one third of the control in the presence of benzamil (Fig. 7B).

**Figure 7.**
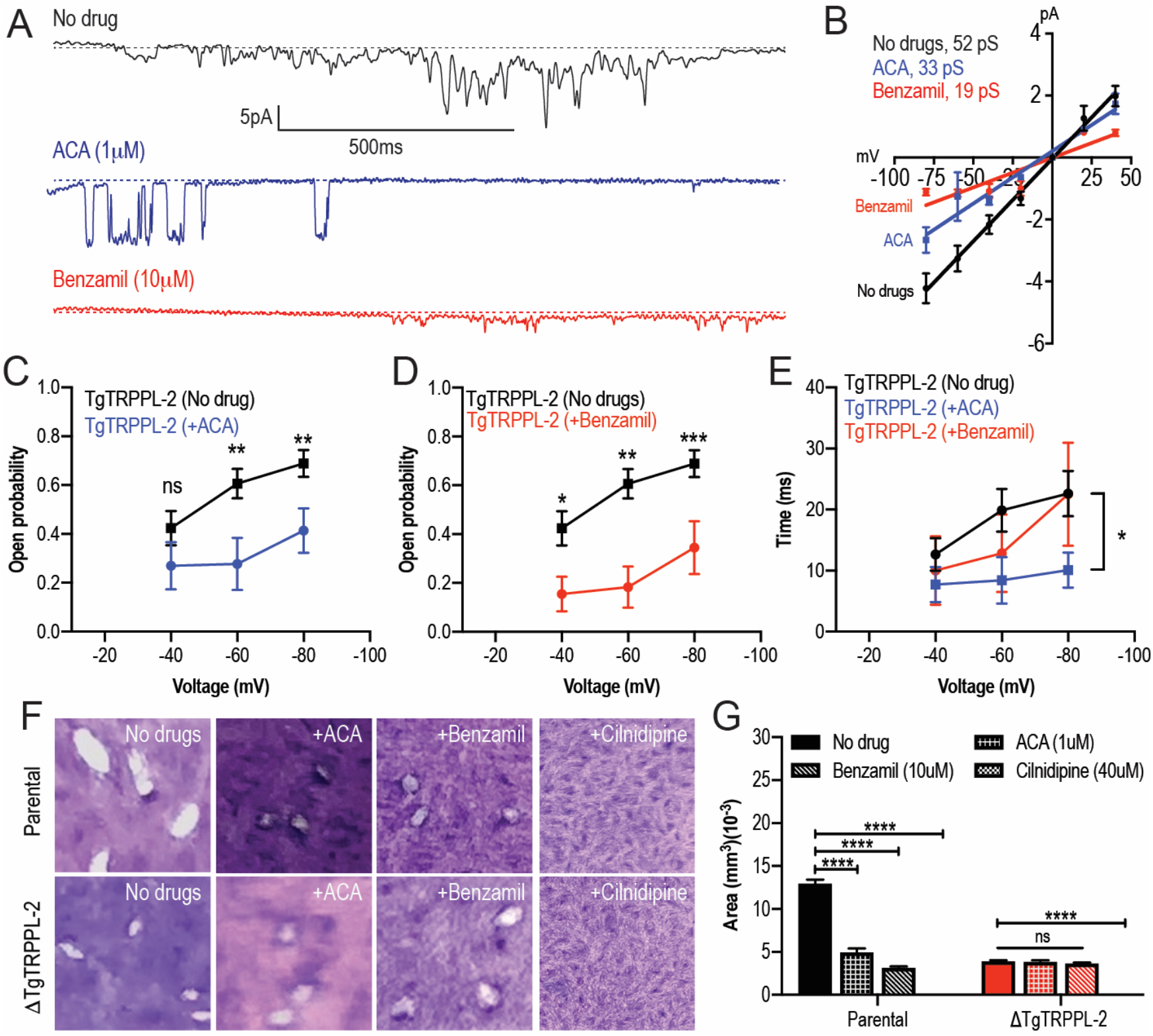
TRP Inhibitors decreased the activity of TgTRPPL-2. **A.** Example of currents recorded of TgTRPPL-2 expressing cells at −80 mV without inhibitors (*Black trace*) compared with the currents in the presence of 1 μM of ACA (*Blue trace*) or 10 μM of benzamil (*Red trace*). **B.** Current-voltage relationship comparing single-channel conductance in the presence of ACA or benzamil. The analysis was done from three independent biological trials and values are means ± SEM. *Inset*, conductance of TgTRPPL-2 in the different conditions analyzed. **C.** Calculated open probability of TgTRPPL-2 expressing cells (*black*) or in the presence of ACA (*blue*). ** *p* <0.006-0.007. **D.** Calculated open probability of TgTRPPL-2 expressing cells (*black*) or in the presence of Benzamil (*red*). Asterisks indicate p values for significance. * *p* < 0.02, ** p< 0.002, *** p< 0.0002. **E.** Average time of channel opening of TgTRPPL-2 expressing cells in the presence of TRP inhibitors. Asterisks indicate p values for significance, * *p* <0.02 **F.** Plaque assay of the *ΔTgTRPPL-2* mutant and the parental strain in the presence of ACA (1 μM), benzamil (10 μM) and cilnidipine (40 μM) after 7 days of growth. **G**. Statistical analysis of plaque sizes done from three independent biological replicates using student’s *t*-test. Values are means ± SEM, **** *p* <0.0001.

Most interesting, we tested both inhibitors, ACA and benzamil in *in vitro* growth assays (Fig. 7F-G, *top panel and parental bars*) and found that they both inhibited *in vitro T. gondii* growth. We calculated the IC_50_ for ACA at 1.4 ± 0.4 μM. Interestingly, neither ACA nor benzamil affected the growth of the *ΔTgTRPPL-2* mutant (Fig. 7G). Note that these cells already grow at a slower rate, which did not change in the presence of the inhibitors. However, cilnidipine, a voltage-gated Ca^2+^ channel blocker, completely inhibited growth of both parental and *ΔTgTRPPL-2* mutant (Fig. 7G).

In conclusion, TgTRPPL-2 is a cation permeable channel that can be inhibited by broad spectrum TRP channel inhibitors. Inhibition of channel activity affects parasite growth.

## DISCUSSION

In this study we report the presence and functional role of a *T. gondii* channel, TgTRPPL-2, that localizes to the plasma membrane and the endoplasmic reticulum. The corresponding gene *TgGT1_310560*, was annotated as hypothetical but was predicted as a transient receptor potential channel based on a bioinformatic analysis of the *T. gondii* genome comparing it with TRP channel genes of mammalian origin [17]. Here, we established that TgTRPPL-2 is important for both Ca^2+^ influx at the PM and Ca^2+^ efflux from the ER of *T. gondii* tachyzoites. TgTRPPL-2, expressed in HEK-3KO cells, conducted currents in high Ca^2+^ solutions and was not voltage dependent. Interestingly, Ca^2+^ itself modulated the conductance of TgTRPPL-2. Broad spectrum TRP channel inhibitors like ACA and benzamil, inhibited the activity of TgTRPPL-2, Ca^2+^ influx in the parasite, as well as parasite growth.

Silencing of TgTRPPL-2 in the *ΔTgTRPPL-2* mutant impacted both invasion and egress of *T. gondii*, resulting in a general growth defect. Extracellular tachyzoites, which are surrounded by high Ca^2+^, are able to use Ca^2+^ influx to stimulate invasion of a new host cell and carry on their lytic cycle. The *ΔTgTRPPL-2* mutant showed a reduction in their host invasion ability suggesting the defect may be due to a reduction in Ca^2+^ influx because of the absence of TgTRPPL-2. Interestingly, the reduction in Ca^2+^ influx (~50%) in the *ΔTgTRPPL-2* mutant was comparable to the reduction of invasion, suggesting that TgTRPPL-2 is involved in the Ca^2+^ influx pathway that stimulates invasion. Delay in the ability of the *ΔTgTRPPL-2* mutant to egress could be caused by a defective efflux of Ca^2+^ from the ER, which was significantly lower in the mutant. This is evidence for the function of TgTRPPL2 as a Ca^2+^ channel at the ER membrane.

The impact of silencing TgTRPPL-2 on *T. gondii* growth, was not total and parasites still were able to perform lytic cycle activities at a reduced rate. The main defects of the *ΔTgTRPPL-2* mutant: invasion, egress, Ca^2+^ influx and ER Ca^2+^ efflux was not complete, likely because more than one mechanism or channel is functional at both locations (PM and ER). We hypothesize the presence of another channel at the PM, likely the one responsible for the Ca^2+^ influx activity that is inhibited by nifedipine [8]. It is also possible that a release channel responsive to IP_3_ may be involved in release of Ca^2+^ from the ER [37] with TgTRPPL-2 having a role in efflux under conditions of ER Ca^2+^ overload.

Numerous observations in *T. gondii* have demonstrated that intracellular Ca^2+^ oscillations in the parasite precede the activation of distinct steps of the lytic cycle [6, 7]. Influx of both extracellular and intracellular Ca^2+^ pools into the parasite cytosol contribute to the activation of downstream signaling pathways decoded into critical biological steps of the parasite lytic cycle [7, 38]. Ca^2+^ influx at the plasma membrane of *T. gondii* is highly regulated, stimulated by cytosolic Ca^2+^ and is operational in extracellular [8] and intracellular replicating tachyzoites [9]. Our data from the *ΔTgTRPPL-2* cells identified TgTRPPL-2 as a functional protein at the plasma membrane and the ER. In these locations it would allow Ca^2+^ influx into the cytosol. The dual localization of TgTRPPL-2 is in accord with other TRP channels in other cells, which showed a dynamic localization between vesicular organelles and the plasma membrane where they facilitate Ca^2+^ influx [39]. In this regard, the mammalian ortholog, PC2, localizes to both the plasma membrane and the ER [40].

*T. gondii* expresses a SERCA-Ca^2+^-ATPase, a P-type ATPase, that couples ATP hydrolysis to the transport of ions across biological membranes (TgSERCA) and localizes to the ER [41]. TgSERCA is sensitive to thapsigargin (thap), a sesquiterpene lactone derived from the plant *Thapsia garganica* [42, 43]. Previous studies showed that inhibition of TgSERCA by thap resulted in cytosolic Ca^2+^ efflux through an unknown channel [8, 36]. In mammalian cells, the passive Ca^2+^ efflux from the ER is thought to prevent ER Ca^2+^ overload and help to maintain the steady-state concentration of luminal Ca^2+^ permitting cytosolic Ca^2+^ signaling [2, 44]. Several membrane proteins have been proposed to be involved in the ER Ca^2+^ efflux/leak pathway including TRP channels [45]. Results from this work support a role for TgTRPPL-2 in ER Ca^2+^ leakage in *T. gondii*. Ca^2+^ efflux from the ER observed after adding thap or Zaprinast was also significantly decreased in the *ΔTgTRPPL-2* mutant. These results support a functional role for TgTRPPL-2 at the membrane of the ER as the constitutive leak channel involved in Ca^2+^ efflux when the store is filled. This could also be the mechanism by which the ER supplies Ca^2+^ to other organelles like the mitochondria or the plant-like vacuole (PLV) a lysosome-like compartment [46].

Previous work from our laboratory showed that Ca^2+^ influx at the plasma membrane did not operate as store-operated Ca^2+^ entry (SOCE) which was shown with experiments testing surrogate ions like Mn^2+^ [8]. This result was supported by the lack of components of the SOCE pathway, STIM and ORAI, in the *T. gondii* genome [17]. However, Ca^2+^ influx was modulated by cytosolic Ca^2+^ [8] and this modulation was absent in the *ΔTgTRPPL-2* mutant supporting a role for TgTRPPL-2 as the channel responsible for Ca^2+^ influx at the PM activated by cytosolic Ca^2+^. TRP channels have been shown to play a role in Ca^2+^-activated-Ca^2+^ entry [47]. Release of Ca^2+^ from intracellular stores like the ER, is also significantly diminished in the *ΔTgTRPPL-2* mutant, which could affect the stimulation of Ca^2+^ influx. However, when using Zaprinast, which raised cytosolic Ca^2+^ at a much higher level than thap, the stimulation of Ca^2+^ influx by cytosolic Ca^2+^ was absent. This further supports that TgTRPPL-2 functions at the PM mediating Ca^2+^ influx and is modulated by cytosolic Ca^2+^.

We showed that TgTRPPL-2 was able to conduct currents with conductance values comparable to the values of mammalian TRP channels [48–50]. Previous work with PC2, showed that Ca^2+^ modulated the activity of PC2 [51–53] [48, 54]. Sustained cytosolic Ca^2+^ increase inhibited PC2 currents [53] while other studies showed that cytosolic Ca^2+^ increase from physiological (100 nM) to μM levels increased the activity of the channel [48, 54]. We observed some of these responses with TgTRPPL-2, as increasing Ca^2+^ inside the pipette (ER luminal) showed a significant decrease in the currents. Comparably, increasing Ca^2+^ concentration in the bath solution (cytosolic) from physiological levels to μM levels showed an increase of 2.5x in the conductance of TgTRPPL-2. Although cytosolic [Ca^2+^] is unlikely to reach those high μM levels, the potential presence of Ca^2+^ microdomains at the plasma membrane or the ER membrane would result in higher concentrations of Ca^2+^ at the exit of the channel due to slow diffusion of Ca^2+^ ions [55–57].

Because PKD channels are cation permeable they could also permeate Na^+^ or K^+^. In the case of TgTRPPL-2 we showed that it can mediate Ca^2+^ transport in the absence of other ions in the solution. We did not determine the ionic selectivity of TgTRPPL-2, and we can only propose that TgTRPPL-2 is a cation permeable channel. In cilia, PKD channels have been described to have relatively high conductance [48, 58]. The conductance calculated for TgTRPPL-2 is within range of what has been described for PC2 in other cells (30-157 pS). However, it is important to note that the properties described for any channel will depend on the experimental approaches used.

Anthranilic acid and benzamil are broad spectrum inhibitors that have the ability to inhibit TRP channel activity. ACA is a weak base that inhibits currents mediated by TRP channels. ACA does not block the pore of the channel as most inhibitors but rather reduce the open probability of the channel. In a similar manner, benzamil is also able to inhibit currents mediated by TRP channels by binding to a site that modulates their activity rather than blocking its pore. In our experiments testing ACA and benzamil we observed that while the inhibitors affected Ca^2+^ influx and growth of the parental cell line, neither affected the already reduced growth and Ca^2+^ influx of the *ΔTgTRPPL-2* mutant. This result combined with the inhibition of TgTRPPL-2 currents impacting both open probability and time that the channel remained open points to TgTRPPL-2 as the target of ACA and benzamil.

Recent studies on Ca^2+^ signaling in *T. gondii* have expanded our understanding of the link between Ca^2+^ and critical facets of parasite biology (i.e., gliding motility, microneme secretion, host cell invasion and egress). However, important molecular players have remained enigmatic, like the PM channels responsible for Ca^2+^ influx and the ER channel responsible for the passive leakage into the cytosol. Characterization of TgTRPPL-2 and its function at the ER and PM fills a small gap in our knowledge of Ca^2+^ signaling and homeostasis in *T. gondii* (Fig. 8). This study is the first biophysical characterization of a channel in *T. gondii* (and any Apicomplexan parasite) and TgTRPPL-2 represents the first identified molecule to mediate Ca^2+^ influx into the cytosol of *T. gondii* at the plasma membrane and the ER. In addition, this study identifies TgTRPPL-2 as a potential target for combatting Toxoplasmosis.

**Figure 8.**
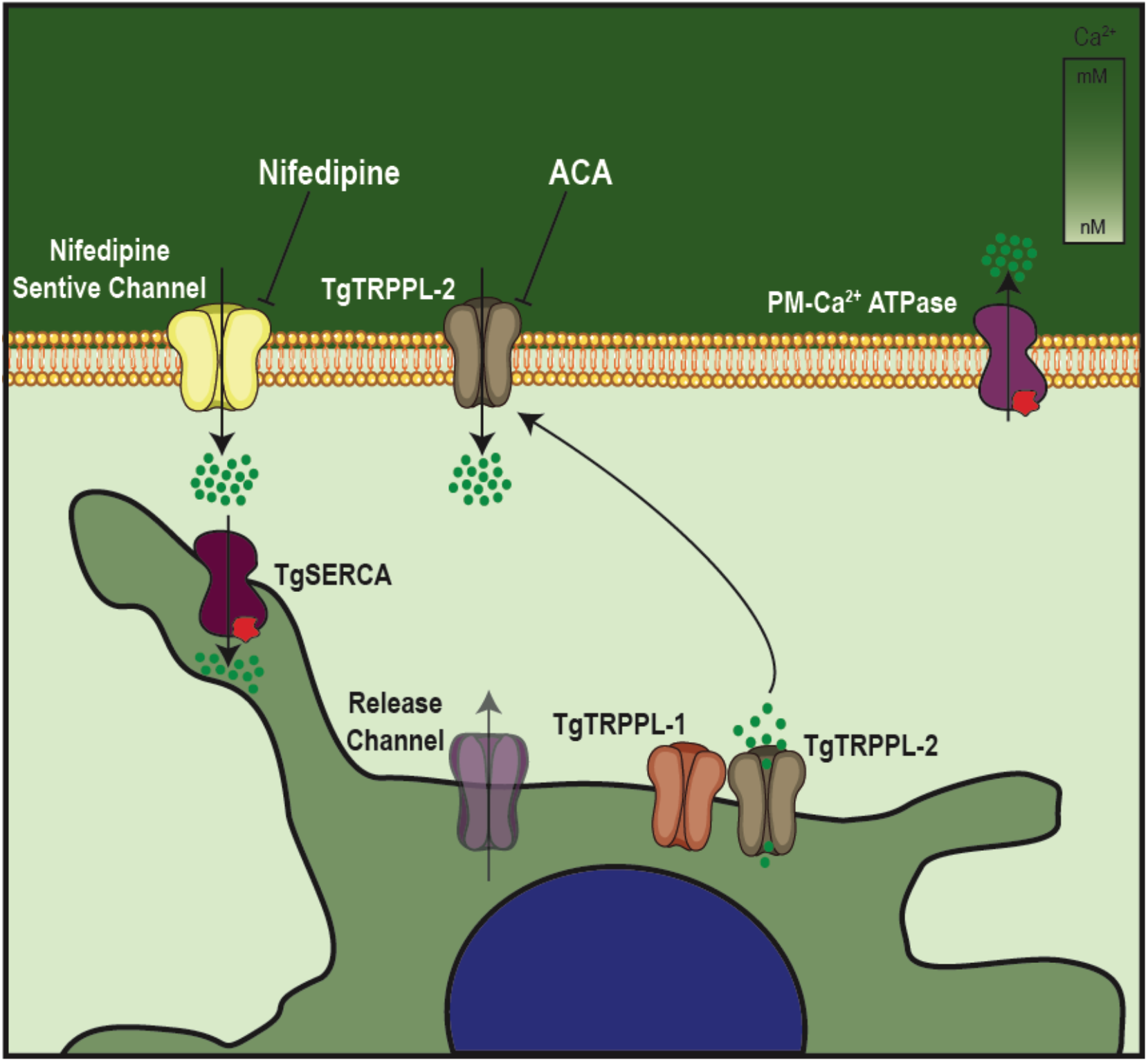
The role of TgTRPPL-2 in Ca^2+^ influx into the cytosol of *T. gondii.* Schematic representation of Ca^2+^ influx into the cytosol of *T. gondii*. Ca^2+^ influx is mediated by two independent Ca^2+^ channels at the PM, a nifedipine-sensitive channel and TgTRPPL-2. TgTRPPL-2 localizes to the plasma membrane as well as the ER. TgTRPPL-2 at the PM is a cation permeable channel that mediates Ca^2+^ influx by a pathway that is activated by high [Ca^2+^] and can be inhibited by broad TRP inhibitors like ACA and benzamil. The presence of the channel at the ER suggest that it may function as a Ca^2+^ efflux channel. Increase in cytosolic [Ca^2+^] can modulate TgTRPPL-2 by allowing the channel to open for longer time thus allowing more Ca^2+^ to enter the cell.

## EXPERIMENTAL PROCEDURES

### *Toxoplasma* growth

All parasite strains were continuously maintained *in vitro* by serial passage in Dulbecco’s modified minimal essential media (DMEM) with 1% FBS, 2.5 *μ*g/ml amphotericin B, 100 *μ*g/ml streptomycin in the human telomerase reverse transcriptase immortalized foreskin fibroblasts (hTERT) [59].

### Generation of mutants

The smHA-LIC-CAT plasmid was used for *in situ* C-terminal tagging of TgTRPPL-2-smHA [19]. Carboxy-terminus tagging was done in the parental line RHTatiΔku80 (TatiΔku80) [23] a parasite line that contains the tetracycline-regulated transactivator system that allows conditional expression of genes [60] and also in which the *ku80* gene was deleted increasing efficiency of homologues recombination [61]. Briefly, a homology region of 974 bp covering the 3’ region of the gene of interest excluding the STOP codon was amplified by PCR using *T. gondii* RH genomic DNA as template and cloned into the plasmid. Plasmids were validated by restriction digest and sequencing. The oligonucleotides primers used for PCR and for creating the gene-tagging plasmids and for PCR validations are listed in Table S3 (Primers T1-T3). Prior to transfection all plasmids were linearized within the region of homology. Approximately 20 μg of plasmid DNA was used for transfection of 1×10^7^ *T. gondii* RHTatiΔKu80 parasites using a Gene Pulser X Cell electroporator (BioRad). Selection was done with 20 μM chloramphenicol, and clones were isolated by limiting dilution. DNA of selected clones were isolated and screened by PCR.

To disrupt the *TgTRPPL-2* (TgGT1_310560) gene a single guide RNA against TgTRPPL-2 was constructed as described [62]. The single guide RNA was mutagenized with the desired sequence in a plasmid that contains the Cas9 using the Q5 Mutagenesis Kit following manufacturer’s instructions. The correct mutation was verified by sequencing. The pyrimethamine-resistant DHFR cassette was amplified by PCR with primers containing 50 bp homology arms of the region upstream and downstream of the start and stop codon of the *TgTRPPL-2* gene. The created sgTgTRPPL-2 CRISPR plasmid was co-transfected with the DHFR cassette (3:1 respectively) into RH tachyzoites. Selection followed with pyrimethamine for 7 days. Parasites were sub-cloned by limiting dilution and screening for clones was done by PCR. The primers used for the creation of the *ΔTgTRPPL-2* are listed in Table S3 (Primer K1-K4).

### Quantitative PCR

Total RNA from parental, *ΔTgTRPPL-2* and *ΔTgTRPPL2-trppl2* parasites was extracted and reversed-transcribed into cDNA. The qPCR reaction was done using the iQ^TM^SYBR Green master mix (BioRad), plus primers, and the reverse-transcribed cDNA (Primers shown in Table S3, Q1-Q2). The qRT-PCR was carried out on a CFX96^TM^ PCR Real-Time detection system (C1000Touch^TM^ Thermal cycler, BioRad). Relative quantification software (CFX Maestro^TM^ software) was used for the analysis and relative expression levels were calculated as the fold change using the formula 2^ΔΔ^CT [63]. Normalization was done using actin and tubulin primers. Experiments were repeated three times with triplicate samples.

### Antibody production of TgTRPPL-2

The antigenic region for TgTRPPL-2 chosen for antibody production was identified using the IEDB suite of antigenicity prediction software. The DNA sequence was amplified from RH genomic DNA and cloned into the pET-32 LIC/EK vector (Novagen), which adds an N-terminal thioredoxin and histidine tag to the expressed protein. Recombinant CP1Ag was expressed and initially purified via a nickel-affinity column (HisPur Thermo Fisher) as previously described [64]. Cleavage of the N-terminal thioredoxin and histidine tag was done by biotinylated thrombin. The antigen was passed again through the nickel column and the purified tag-less antigen was gently eluted using 10 mM imidazole. Antibodies in mice were generated as previously [65]. CD1 mice (Charles River) were inoculated intraperitoneally with 100 μg of the TgTRPPL-2 peptide mixed with complete Freund's adjuvant, followed by two boosts with 50 μg of the TgTRPPL-2 peptide in incomplete Freund's adjuvant. The final serum was collected by cardiac puncture after CO_2_ euthanasia. We created a αSERCA antibody for co-localization studies of the TgTRPPL-2. The phosphorylation (P) and nucleotide binding (N) domains of *Tg*SERCA were cloned into XmaI and HindIII sites of pQE-80L plasmid for expression in *Escherichia coli* BL21-CodonPlus competent cells. Purified antigen was used to immunize Guinea pigs with 0.2 mg of antigen mixed with Freund’s Complete Adjuvant, followed by two boosts of 0.1 mg antigen mixed with Freund’s Incomplete Adjuvant (Sigma F5506). The resulting antibodies were used at 1:1,000 for western blots. The animal protocol used was approved by the UGA Institutional Animal Care and Use Committee (IACUC).

### Western blot analysis

SDS-polyacrylamide gel electrophoresis (SDS-PAGE) followed established protocols [66]. Lysates were prepared by resuspending a pellet of 1 × 10^8^ tachyzoites in 50 μL of Cell Lytic^M^ lysis buffer containing 12.5 U benzonase and 1 × protease cocktail inhibitor (P8340 Sigma). The reaction was stopped with one volume of 2% SDS and 1 mM EDTA. Total lysates were boiled in Laemmli sample buffer (BioRad). Immunoblotting followed established protocols using mouse anti-HA monoclonal antibody (1:1,000) (Roche). Detection was done using the Odyssey Clx LICOR system using goat anti-mouse IRDye800WC (1:10,000). Loading control for westerns were done with primary mouse-anti-tubulin antibodies at a 1:15,000 dilution and goat anti-mouse IRDDye800WC as secondary (1:10,000).

### Immunofluorescence microscopy

Extracellular parasites were collected and purified as previously [67]. Parasites were washed once with buffer A with glucose (BAG, 116 mM NaCl, 5.4 mM KCl, 0.8 mM MgSO_4_, 5.5 mM glucose and 50 mM HEPES, pH 7.4) and an aliquot of 2 × 10^4^ parasites was overlaid on a coverslip previously treated with poly-L-Lysine. Intracellular tachyzoites were grown on hTERT cells on coverslips. Both extracellular and intracellular parasites were fixed with 3% paraformaldehyde for 20 min at room temperature (RT), permeabilized with 0.3% Triton X-100, blocked with 3% bovine serum albumin (BSA), and exposed to primary antibodies (Ratα-HA 1:100). The secondary antibodies used were goat-αrat Alexa Fluor 488 (Life Technologies) at a 1:1,000 dilution. For co-localization studies we used α-Sag1 (1:1,000) as membrane marker and α-TgSERCA as ER marker (1:1,000). Slides were examined using an Olympus IX-71 inverted fluorescence microscope with a photometric CoolSNAP HQ charge-coupled device (CCD) camera driven by DeltaVision software (Applied Precision, Seattle, WA).

### Immunoprecipitation assays

Freshly lysed tachyzoites expressing TgTRPPL-2-smHA were collected and filtered through an 8 *μ*M membrane (Whatman). Tachyzoites were washed twice in BAG and resuspended in lysis buffer (50mM Tris-HCl, pH 7.4, 150 mM KCl, 1 mM EDTA, 0.4% NP-40) to a final concentration of 2 × 10^9^ total cells. Lysis was allowed to proceed for thirty minutes at 4°C and cells were centrifuged at 15,000 × g for 20 min. Immunoprecipitation of TgTRPPL-2-smHA protein was performed using the Pierce HA Tag/Co-IP Kit (Thermo Fisher Scientific, Waltham, MA) according to manufacturer’s instructions. Briefly, HA magnetic beads were washed twice in lysis buffer and mixed with the parasite lysate by vortexing for 1 h at RT. Beads were collected and the flow-through fraction was saved for further analysis. Beads were washed twice in wash buffer (50mM Tris-HCl, pH 7.4, 150 mM KCl, 1 mM EDTA, 0.1% NP-40) and once in ddH_2_O by gentle mixing. The tagged protein was recovered by mixing the beads with 1x Laemmli buffer and heated at 65°C for 10 min. The supernatant was collected and used for PAGE and western blots. The corresponding band was cut and resuspended in water and analyzed using LC-Mass Spectrometry. Samples were sent to the Proteomics and Mass Spectrometry Core Facility at the University of Georgia for analysis. The average counts that were obtained from two biological samples are shown in Table S2. Proteins with counts higher than 3 are shown.

### Growth and Invasion Assays

Plaque assays were done as previously described, with slight modifications [67]. Briefly, 200 egressed tachyzoites were allowed to infect confluent hTERT cells for 7 days. After seven days cells were fixed with ethanol and stained with crystal violet. Plaque sizes were analyzed using FIJI [68]. Invasion assay were performed as previously described, with slight modifications [28]. A subconfluent monolayer of HFF cells were infected with 2×10^7^ tachyzoites in the presence of 1.8 mM or 0.5 mM Ca^2+^ and placed for 20 min on ice and subsequently transferred for 5 min to a 37°C water bath for parasite invasion. Cells were immediately fixed with 3% paraformaldehyde for 20 min. Extracellular parasites (attached) were stained using RabbitαSag1 (1:1,000) prior to permeabilization while intracellular parasites (invaded) were stained with MouseαSag1 (1:200). Secondary antibodies were goat-αrabbit Alexa Fluor 546 (1:1,000) and goat-αmouse Alexa Fluor 488 (1:1,000). Images were taken with an Olympus IX-71 inverted fluorescence microscope with a Photometric Cool SNAP HQ CCD camera driven by DeltaVision software (Applied Precision, Seattle, WA). Quantification was made of ten-fields of view at a 1000 magnification from three independent biological replicates. Percentage of invaded vs attached was quantified by dividing the number of parasites invaded or attached by the total parasites quantified in the field of view.

### Egress experiments

hTERT cells were infected with 5×10^5^ of RH or *ΔTgTRPPL-2* tachyzoites. 24 h after infection parasitophorous vacuoles were observed by microscopy and washed with Ringer’s buffer (155 mM NaCl, 3 mM KCl, 1 mM MgCl2, 3 mM NaH_2_PO_4_H_2_O, 10 mM HEPES, pH 7.3, and 5 mM glucose). Ringer’s buffer was used as extracellular buffer in the presence or absence of 1.8 mM Ca^2+^. Drugs were added in Ringer’s buffer 30 sec after imaging at the following concentrations: saponin (0.02%) or Zaprinast (100 μM). Images were acquired in a time-lapse mode with an acquisition rate of 3 sec for 12-20 min For statistical analysis, egress time was quantified as the first parasite to egress out of the parasitophorous vacuole. Statistical analysis was done for 3 independent biological replicates and at least 5 PVs per experiment.

### Cytosolic Ca^2+^ measurements

Parasites were loaded with Fura2-AM as described in [36]. Briefly, fresh lysed parasites were washed twice at 1,800 rpm for 10 min at room temperature in buffer A (BAG) (116 mM NaCl, 5.4 mM KCl, 0.8 mM MgSO4, 5.5 mM d-glucose and 50 mM Hepes, pH 7.4). Parasites were resuspended to a final density of 1×10^9^ parasites/mL in loading buffer (Ringer’s plus 1.5% sucrose, and 5 μM Fura2-AM). The suspension was incubated for 26 min at 26 °C with mild agitation. Subsequently, the parasites were washed twice with Ringer’s buffer to remove extracellular dye. Parasites were resuspended to a final density of 1×10^9^ parasites/mL in Ringer’s buffer and kept in ice. For fluorescence measurements, 2×10^7^ parasites/mL were placed in a cuvette with 2.5 mL of Ringer’s buffer. The cuvette was placed in a thermostatically controlled Hitachi F-7000 fluorescence spectrophotometer. Excitation was at 340 and 380 nm, and emission at 510 nm. The Fura2-AM fluorescence relationship to intracellular Ca^2+^ concentration ([Ca^2+^]_i_) was calibrated from the ratio of 340/380 nm fluorescence values after subtraction of the background fluorescence of the cells at 340 and 380 nm as previously described [69]. Changes in [Ca^2+^]_i_ (ΔF [Ca^2+^]) were measured by subtracting the highest peak of Ca^2+^ in the first 20 s after addition of Ca^2+^ or 100 s after the addition of drugs minus the baseline.

### Cell transfections and culture of HEK-3KO Cells

Total RNA of wild type *T. gondii* were extracted and reversed transcribed into cDNA. TgTRPPL-2 whole cDNA was amplified using primers shown in Table S3 (Primers C1-C6). The amplified cDNA was cloned into the Zero Blunt TOPO vector using the cloning kit per manufacturers instruction. Correct insertion was verified by colony PCR using M13F and M13R primers. Restriction digests was performed to remove the insert from the vector using the following restriction enzymes: BamHI and AvrII. The purified *TgTRPPL-2 cDNA* was ligated to linearized pCDNA 3.1 plasmid. Ligation to the vector was confirmed by PCR and sequencing. Purified TRPPL-2-pCDNA was used to co-transfect DT40-3KO cells.

HEK cells which have the 3 endogenous isoforms of the IP_3_ receptor knocked out were a gift from Dr. David Yule [31, 70]. The cells were maintained in Dulbecco’s modified minimal essential media (DMEM) with 10% fetal bovine serum 2.5 *μ*g/ml amphotericin B and 100 *μ*g/ml streptomycin. Cells were transiently transfected as previously described [71] with 2.5 μg of TgTRPPL-2, PC2 or RFP DNA targeted to the ER. Each plasmid DNA were diluted in 200 μL of Opti-MEM with 25 μL of polyethylenimine and incubated for 10 min. The mix was then added to semi confluent HEK-3KO cells in a dropwise manner, and 24 h later the media was changed.

### Preparation of nuclear extracts

48 hours after transfection, cells were collected and the nucleus extracted as previously described [72]. 2×10^7^ of transiently transfected cells were collected in ice cold PBS. Cells were spun down and washed twice in PBS and resuspended in Nuclei Isolation Solution (150 mM KCl, 250 mM Sucrose, 10 mM Tris-HCl, 1.4 mM β-mercaptoethanol, 0.2 mM PMSF, pH 7.3). Cells were homogenized with a homogenizer and stored on ice. 100 μL of nuclei were transferred to cover slips previously coated with poly-L-lysine and incubated for 20 minutes before filling the chamber with bath solution.

### Patch clamp of nuclear membranes

Nuclear extract expressing TgTRPPL-2 or the control gCaMPER [73] were used for analysis. Electrical currents were recorded using Standard Wall Borosilicate Capillaries (Harvard Bioscience, Massachusetts) with 10-15 MΩ resistance. Holding potentials were maintained at 0 mV. The internal solution contained: 140 mM KCl or CsCl, 10 mM HEPES, 1.8 mM or 10 mM free Ca^2+^ adjusted with EGTA. Standard Bath Solution contained the same reagents and concentrations as the pipette solution using 100 nM of free Ca^2+^. The single-channel conductance was obtained from the current-voltage relationship for each condition tested. Ca^2+^ currents were elicited by applying pulses from −80 mV up to 20 mV for 25 seconds. Analysis of amplitude, open probability and channel conductance were done using a 45 kHz filter. Data for recording was collected using the HEKA Electronic Patch Clamp EPC10 (Harvard Bioscience, Massachusetts).

### Statistics

Statistical analyses were performed by Student’s *t-test* using GraphPad PRISM version 8.2. All experimental data were analyzed from at least three independent biological replicates. Error bars shown represent standard error of the mean (SEM) of the biological replicates analyzed. For the electrophysiological analysis a total of 3 cells per biological replicate (9 total cells) were analyzed. Each cell was depolarized a total of 5 times per experimental conditions.

## Acknowledgements

We thank Drs. David Yule for providing the HEK-3KO cells for electrophysiological analyses, John Boothroyd for antibodies against SAG1 and Boris Striepen for the Cosmid for complementation. Catherine Li prepared the antigen used to generate the anti-SERCA antibody. The super resolution microscope is part of the Biomedical Microscopy Core (BMC) of the University of Georgia. We thank the University of Georgia Graduate School for a Summer Research Travel Grant to KMMN. This work was supported by the U.S. National Institutes of Health grant AI128356 to SNJM and R00 DK101585 to IYK. KMMN was partially supported through a fellowship funded by a T32 training grant, 5T32AI060546.

## Author Contributions

**KMMN**, performed and coordinated most of the experiments, analyzed the data, wrote the manuscript; **NC**, developed the specific antibody and contributed to its validation; **MAHT**, developed the knockout strain and its validation; **IK**, writing, review, editing, analysis and interpretation of data; **SNJM**, coordinated the project and experiments, contributed resources and wrote the manuscript.

## Notes

### Competing Interest Statement

The authors have declared no competing interest.

### Summary of Updates

Name of authors, abstract and description of the mutant was revised

