## Supplementary material for "Calcium signaling through a Transient Receptor Channel is important for *Toxoplasma gondii* growth": Supplemetal tables and figures

**Márquez-Nogueras et al**

**Supplemental tables and Figures**

**Table S1:** Top 10 hits of HHPRED analysis of TgTRPPL-2.

| Name | Probability | E-value | AA lenght | PDB Reference |
|---|---|---|---|---|
| **Polycystic kidney disease 2-like 1** | 99.45 | 7.10 $E^{-11}$ | 805 | 6DU8_A |
| **Polycystin-2; PKD2** | 99.44 | 1.10 $E^{-10}$ | 756 | 6WB8_D |
| **Polycystic kidney disease 2-like 1** | 99.39 | 2.10 $E^{-10}$ | 566 | 5Z1W_C |
| **Polycystin-2, Polycystin-1** | 99.38 | 2.40 $E^{-10}$ | 577 | 6A70_A |
| **Polycystin-2** | 99.35 | 2.40 $E^{-10}$ | 968 | 5MKE_A |
| **Polycystin-2, Polycystin-1** | 99.3 | 2.80 $E^{-10}$ | 1153 | 6A70_B |
| **Polycystin-2; TRP channel, PKD2** | 99.22 | 3.30 $E^{-09}$ | 510 | 5T4D_A |
| **TRPV2; Transport protein, TRP channel** | 98.52 | 6.6 $E^{-05}$ | 613 | 5AN8_B |
| **Mucolipin-3; TRP channel, lysosomal** | 98.31 | 0.00031 | 558 | 6AYF_C |
| **Transient receptor potential cation channel** | 98.28 | 0.00015 | 639 | 6LGP_D |

**Table S2:** List of Mass spectrometry hits by TgTRPPL-2 Immunoprecipitation*.

| Gene ID | Description | Phenotype# | TMD | Average counts |
|---|---|---|---|---|
| TgGT1_310560 | Hypothetical protein (TgTRPPL-2) | -2.49 | 13 | 3 |
| TgGT1_247370 | Hypothetical protein (TgTRPPL-1) | -1.42 | 13 | 3 |
| TgGT1_214300 | Hypothetical Protein | -0.36 | 9 | 3 |
| TgGT1_280560 | Selenide, water dikinase | 0.07 | 0 | 6 |
| TgGT1_201680 | Putative eukaryotic initiation factor-3 subunit 10 | -4.55 | 0 | 6 |
| TgGT1_228170 | Inner membrane complex protein IMC2A | -3.28 | 1 | 5.5 |
| TgGT1_212300 | Hypothetical protein | 0.69 | 1 | 3 |
| TgGT1_229180 | HEAT repeat-containing protein | -5.05 | 0 | 3 |

*See Experimental Procedures for the protocol.

### Fitness score for each gene was obtained from ToxoDB.

**Table S3:** Primers used in this work

| | | |
|---|---|---|
| ***Endogenous Tagging of TgTRPPL2*** | | |
| **T1** | TgTRPPL2_pLic_F | TACTTCCAATCCAATTTAATGCGAGAAGCGCATT GAGGAATGG |
| **T2** | TgTRPPL2_pLic_F | TCCTCCACTTCCAATTTTAGCCTCTTCTCCCAGG ATGTTGACGC |
| **T3** | TgTRPPL2_Validation_Tag_F | TATGTGTGCCTGCCTGCGCAT |
| ***Disruption of TgTRPPL2*** | | |
| **K1** | TgTRPPL2_Cas9_gRNA_F | TATGTCACATGTCTTTTCTCGTTTTAGAGCTAGAA ATAGCAAG |
| **K2** | TgTRPPL2_DHFR_F | CTTTGGTTTCCCTCTCTCGTCCATGAAGCTTCGCC AGGCTGTAAATCC |
| **K3** | TgTRPPL2_DHFR_R | TGGACGCCCAGCTCGACATGTCATCCTGCAAGTG CATAGAAGGA |
| **K4** | TgTRPPL2_Validation R | CGATGAGGTGGATGTAGCTGAATG |
| ***RT-PCR of TgTRPPL2*** | | |
| **Q1** | TgTRPPL2_qPCR_F | GAGCTCCGACGCAGGCCAGCAG |
| **Q2** | TgTRPPL2_qPCR_R | CCCGGGCGATGAGGTGGATGTAGCTGAATG |
| ***Cloning for Heterologous expression in DT-40-3KO cells*** | | |
| **C1** | pCDNA3_TgTRPPL2_F | cagatatccatcacactggcATGCATGCATTCGACGAC |
| **C2** | tdTomato_TgTRPPL2_R | tgctcaccatCTCTTCTCCCAGGATGTTG |
| **C3** | TgTRPPL2_tdTomato_F | gggagaagagATGGTGAGCAAGGGCGAG |
| **C4** | pCDNA3_TgTRPPL2_tdTomato _R | acactatagaatagggccctCTACTTGTACAGCTCGTCC ATG |
| **C5** | TgTRPPL2_Validation_F | GCAAGAAGAAGAAACGACGCAAG |
| **C6** | TgTRPPL2_Validation_R | CTTTGAGGTCCTAGTTCACCTCCGA |

**Table S4.** Composition of the solutions used for the electrophysiological analysis.

| Reagents | Concentration (mM) | | | | |
|---|---|---|---|---|---|
| | **Solution A** | **Solution B** | **Solution C** | **Solution D** | **Solution E** |
| KCl | 140 | 140 | 140 | - | - |
| CsCl | - | - | - | 140 | 140 |
| EGTA | 0.1 | 0.1 | 0.1 | 0.1 | 0.1 |
| Intracellular $Ca^{2+}$ | 1.8 | 10 | 1.8 | 1.8 | 10 |
| Extracellular $Ca^{2+}$ | 0.0001 | 0.0001 | 0.01 | 0.0001 | 0.0001 |

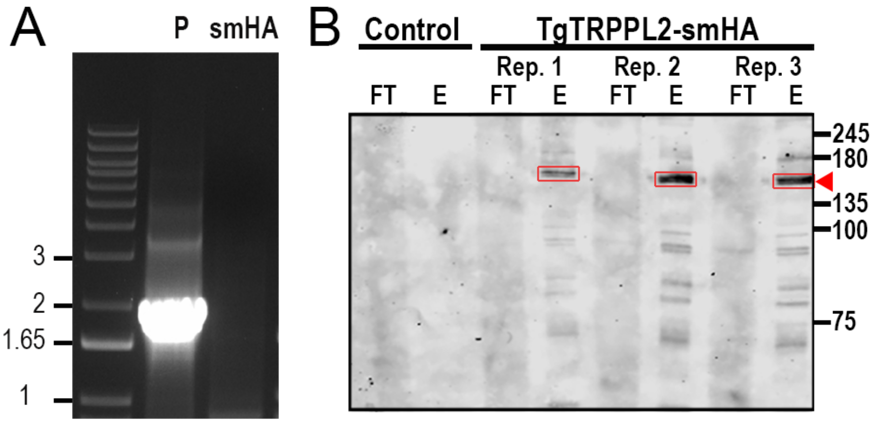


**Figure S1. Validation of C-terminal tagging of TgTRPPL-2-smHA. A.** Amplification of ~2 kB in the TgTRPPL-2-smHA cell line validating the correct integration of TgTRPPL-2-smHA. **B.** Western blot of three biological replicates of immunoprecipitated TgTRPPL-2-smHA using αHA antibody (1:1,000) shows a band of ~150kDa (highlighted by *red boxes* and *arrow*). No band is present in the control cell line.

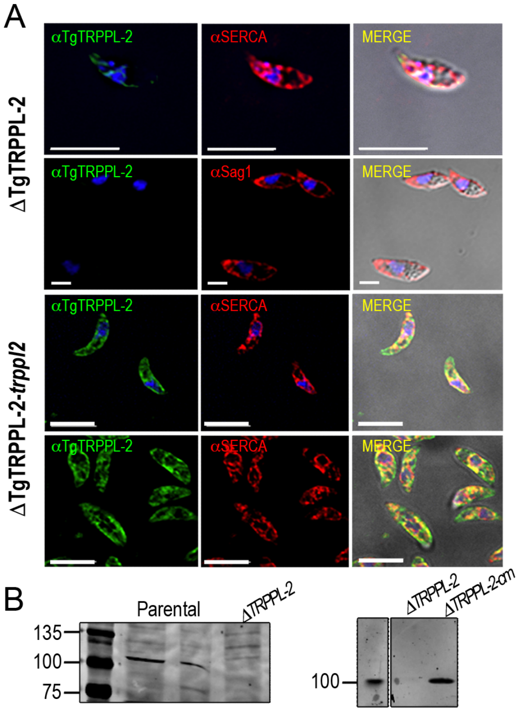


**Figure S2. Validation of the anti-TgTRPPL-2 antibody. A,** IFAs of extracellular tachyzoites with αTgTRPPL-2 (1:1,000) co-localized with αSERCA (1:1,000) in ΔTgTRPPL-2 and ΔTgTRPPL-2-*trppl2*. Images were taken with equivalent time and laser power. B, Western blots of lysates from parental, *ΔTgTRPPL-2* and ΔTgTRPPL-2-*trppl2* cells were run in an SDS-PAGE cell and developed with the anti-TgTRPPL-2 antibody at 1;1,000.

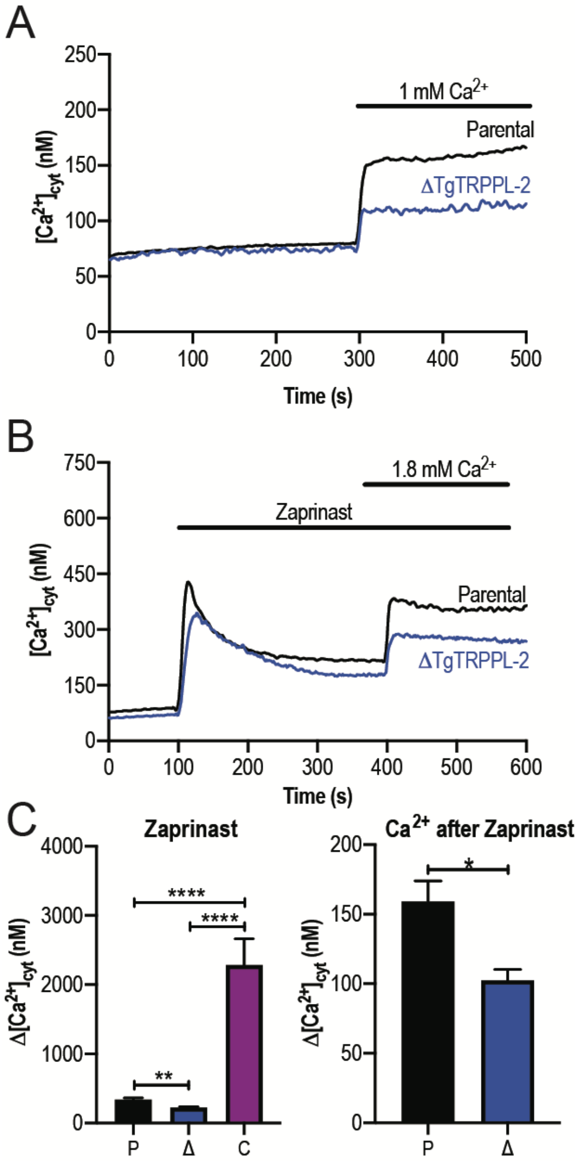


**Figure S3. TgTRPPL-2 regulates $Ca^{2+}$ in *T. gondii*.** **A.** Cytosolic $Ca^{2+}$ measurements of Fura-2 loaded tachyzoites of the parental and *ΔTgTRPPL-2* mutant after addition of 1 mM extracellular $Ca^{2+}$ at 300 s. **B.** Cytosolic $Ca^{2+}$ measurement after the addition of Zaprinast (100 $\mu$M) at 100 s and $Ca^{2+}$ influx was stimulated by the addition of 1.8 mM extracellular $Ca^{2+}$ at 400s. **C.** Change in cytosolic $Ca^{2+}$ 15 s after the addition of Zaprinast *(Labeled Zaprinast)* and 20 s after the addition of 1.8 mM of extracellular $Ca^{2+}$ *(Labeled $Ca^{2+}$ after Zaprinast)*. Values are means ± SEM, ** $p < 0.007$, **** $p < 0.0001$.

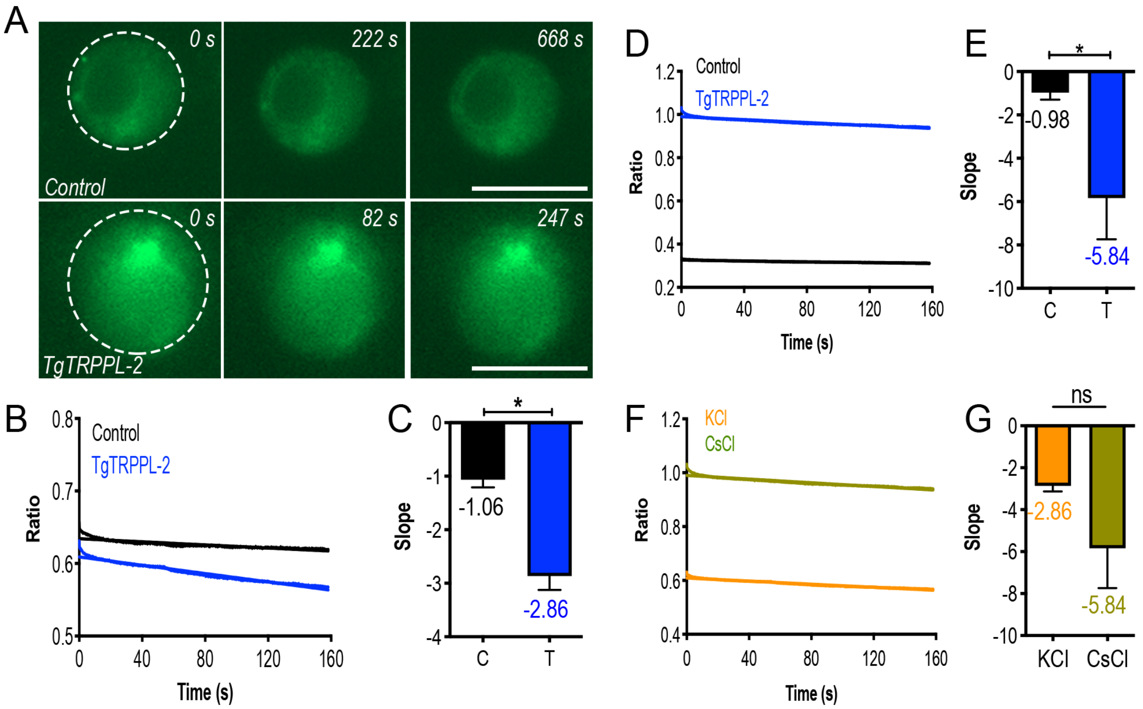


**Figure S4. Measurement of ER Calcium of DT40-3KO cells expressing TgTRPPL-2. A.** Fluorescence of ER calcium in ER-RFP-DT40 (*Control*) vs. TgTRPPL-2-DT40 (*TgTRPPL-2*) in a high calcium-potassium solution of patched nuclear membranes while the membrane is depolarized. **B.** Quantification of fluorescence of patched cells while the membrane is depolarized from -80 to +20 mV in a high calcium-potassium solution. **C.** Quantification of the slope of fluorescence in *B* comparing TgTRPPL-2-DT40-3KO cells versus the control cells. Slope of the fluorescence was quantified based on the 5 technical replicates of the artificial membrane depolarization in each cell analyzed. Asterisk indicate p value for significant difference. Values represent the mean of the slope shown in *B*. * $p < 0.01$. **D.** Quantification of fluorescence of patched cells while the membrane is depolarized from -80 to +20 mV in a High calcium-cesium solution. **E.** Quantification of the slope of fluorescence in *D* comparing TgTRPPL-2-DT40-3KO cells versus control cells. The slope was quantified for the five technical replicates of patched membranes. Asterisk indicate p value for significant difference. Values represent the mean of the slope shown in *D*. * $p < 0.03$. **F.** Comparison of the fluorescence of patched nuclear extract of TgTRPPL-2-DT40 cells while the membrane was artificially depolarized. **G.** Quantification of the slope of fluorescence of patched TgTRPPL-2-DT40-3KO nuclear extracts in different experimental conditions. Slope of the fluorescence was quantified based on the 5 technical replicates of the artificial membrane depolarization in each cell analyzed.
